# The *Endo-GeneScreen* Platform Identifies Drug-Like Probes that Regulate Endogenous Protein Levels within Physiological Contexts

**DOI:** 10.1101/2025.03.13.643156

**Authors:** Preston Samowitz, Laszlo Radnai, Thomas Vaissiere, Sheldon D. Michaelson, Camilo Rojas, Ryan Mitchell, Murat Kilinc, Austin Edwards, Justin Shumate, Richard Hawkins, Virneliz Fernandez-Vega, Timothy P. Spicer, Louis Scampavia, Theodore Kamenecka, Courtney A. Miller, Gavin Rumbaugh

## Abstract

Traditional phenotypic drug discovery platforms have suffered from poor scalability and a lack of mechanistic understanding of newly discovered phenotypic probes. To address this, we created Endo-*GeneScreen* (EGS), a high-throughput enabled screening platform that identifies bioactive small molecules capable of regulating endogenous protein expression encoded by any preselected target gene within a biologically appropriate context. As a proof-of-concept, *EGS* successfully identified drug candidates that up-regulate endogenous expression of neuronal *Syngap1,* a gene that causes a neurodevelopmental disorder when haploinsufficient. For example, SR-1815, a previously unknown and undescribed kinase inhibitor, alleviated major cellular consequences of *Syngap1* loss-of-function by restoring normal SynGAP protein levels and dampening neuronal hyperactivity within haploinsufficient neurons. Moreover, we demonstrate that *EGS* assays accelerate preclinical development of identified drug candidates and facilitate mode-of-action deconvolution studies. Thus, *EGS* identifies first-in-class bioactive small molecule probes that promote biological discovery and precision therapeutic development.

## INTRODUCTION

High-throughput phenotypic screening (HTS) has re-emerged as a powerful approach in drug discovery and functional genomics^1–3^. Unlike target-based strategies that begin with a predefined molecular target, phenotypic screening starts with an observable biological outcome (phenotype) and then identifies compounds or genetic perturbations that produce the desired effect. This forward pharmacology approach allows for the unbiased discovery of bioactive agents, often capturing complex, system-level responses that target-centric methods might overlook. As a result, phenotypic strategies have been credited with the discovery of many first-in-class drug candidates that regulate essential cell signaling pathways implicated in disease ^4–6^.

Phenotypic assays that measure expression of proteins encoded by critical genes required for cellular health promotes advancements in drug development and discovery biology^7^. By measuring changes in a protein’s native expression level, it is possible to identify drug-like probes that increase (up-regulate)^8^ or decrease (down-regulate)^9^ their abundance, thereby expanding utility to discover both activators and inhibitors of disease-relevant pathways. Performing assays in native cellular models further improves translational relevance, as hits are identified within a context that closely mirrors relevant pathobiology. By embracing this unbiased, context-rich strategy, one can reveal novel regulatory pathways and therapeutic strategies that would likely remain hidden in target-centric approaches. Traditionally, there has been a significant trade-off between biological appropriateness and scalability (e.g., how many agents can be screened) – as the context becomes *<u>more</u>* relevant, *<u>fewer</u>*compounds can be screened because the physiologically relevant environment is typically more complex. Moreover, phenotypic screening in general suffers from a lack of target-based information with respect to how a cellular phenotype is altered in response to probe application. Together, these factors have limited the application of phenotypic screening to disease biology and drug development, particularly for disorders of the brain.

We reasoned that recent advances in biotechnology can be leveraged to develop phenotypic screening platforms that increase the scale of screening campaigns without sacrificing the appropriateness of the biological context. Additional advances in chemical biology and chemo-proteomics have also improved the success rate of molecular target deconvolution of phenotypic probes^10^. Here, we demonstrate how advances in endogenous protein detection within native cells^11^, lab automation, and the recent widespread availability of massive, commercially available libraries of drug-like bioactive molecules were integrated to create the *Endo*-GeneScreen (*EGS*) platform **(Fig. 1)**. This flexible HTS-enabled screening platform identifies small molecules capable of regulating endogenous protein expression encoded from any preselected target gene within biologically appropriate cellular contexts. We focused on small molecule screening because these agents remain the gold standard therapeutic for most disease indications^12^, and their chemical diversity can reveal previously unknown druggable pockets in nucleic acids and proteins^13^, a process that facilitates biological discovery.

**Figure 1.**
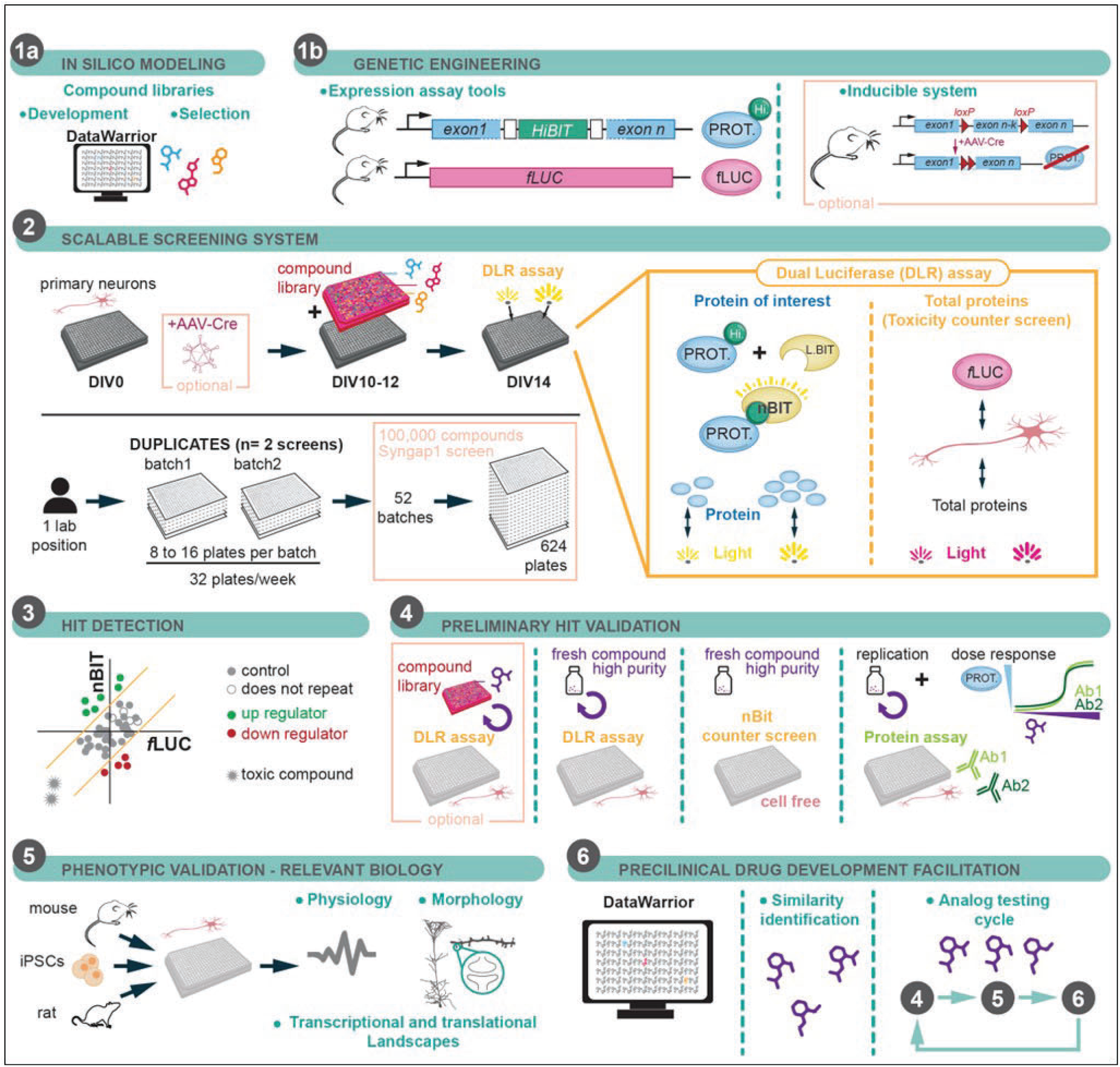
Overview of *Endo*-GeneScreen (EGS) Platform. 1a) A relevant screening library is selected based on the cellular context of interest (e.g. a CNS-targeted library for brain cells; only relevant if drug-development is a future goal). **1b)** mouse models are developed; FLAG-HiBIT tag is inserted into gene of interest. This tag could be inserted into a transgene if protein lowering in a cellular context is the goal. Inclusion of a cKO mouse model is a must if protein upregulation is desired within a gene loss-of-function cellular context. **2)** DLR assay workflow showing how endogenous protein are detected within a specific cellular context; screening plates required for 100k compound screen for screening within cortical mouse neurons. **3)** After completion of the screen, data is visualized on a per-plate basis and hit detection and filtering algorithms are implemented to identify compounds of interest. **4)** Preliminary hits are run through a biological validation workflow that focuses on replication and repeatability ultimately relying on protein detection using KO-validated antibodies (if available). **5)** Probes that are validated at the protein level within the cellular context of interest are then tested in a series of low-throughput phenotypic assays that ideally are known to be modified by altered expression of the protein of interest. **6)** Validated probes that also regulate cell-context-specific phenotypes are then explored at the chemical level by obtaining ∼100 closely related analogues. This determines the extent to which a chemical series with promise can be modified by medicinal chemistry.

As a proof-of-concept, we demonstrate how each component of the platform is used to identify, validate, develop, and define the mode-of-action of drug-like small molecules that upregulate expression of the target gene, *SYNGAP1/Syngap1* (*HUMAN/Mouse*; *Mouse* from now on). *Syngap1* was selected for multiple reasons. First, *de novo* mutations in this gene that lead to haploinsufficiency are one of the most common genetic causes of sporadic neurodevelopmental disorders associated with intellectual disability, autism, and epilepsy^14–18^. Thus, among many potential uses, this platform can be used to screen for “boosters” of autosomal dominant (AutD) haploinsufficiency genes. These genes cause disease through *de novo* mutations that lead to reduced functional protein expression^19, 20^. As such, small molecule “boosters” would, in principle, address the root cause of these genetic conditions. Second, this gene principally functions in differentiated cortical excitatory neurons^21, 22^, a notoriously difficult cellular system to work with at scale^23^. Third, excellent *in vitro* and *in vivo* models for *Syngap1* haploinsufficiency have been developed and extensively validated^24–26^. These models provide a vertically integrated ecosystem that can be used to first screen for probes that upregulate SynGAP protein in an appropriate context (e.g., cortical neurons) and then eventually validate the effectiveness of these probes in the same system. Using a *Syngap1*-focused variation of the EGS platform, we identified SR-1815, a first-in-class small molecule that restored low SynGAP expression and mitigated synaptic and neuronal hyperexcitability, three major cellular consequences of *Syngap1* haploinsufficiency. Moreover, we demonstrate how *EGS* workflows accelerate preclinical drug development by jump-starting medicinal chemistry-based improvement of identified compounds. Finally, we show that *EGS* assays, when combined with emerging chemical biology and molecular genetic approaches, successfully deconvoluted the key molecular targets and mode-of-action of SR-1815, a novel kinase inhibitor that regulates splicing relevant to neurological disorders and cancer (*see companion paper -* Douglas et al., Biorxiv, 2025).

## RESULTS

### Development of EGS scalable assays: Tracking endogenous target protein expression within disease-appropriate cellular contexts

*EGS* is built upon a series of phenotypic assays that report relative changes in endogenous protein encoded by a pre-selected target gene within a biologically-relevant cellular context. The foundational principle of the platform concept was to develop scalable endogenous protein expression assays within relevant disease-modeling cellular contexts, such as 2D/3D cultures from primary cells derived from animal models or induced cellular models derived from patient iPSCs **(Fig. 1)**. When engineering the scalable assays, we set several design parameters: 1) *<u>flexible targeting</u>* - any gene of interest can be targeted with the assays; 2) *<u>relevant cellular context</u>* – assay must be carried out in disease-modeling cellular systems; 3) *<u>sufficient scalability</u>* – HTS- capable to enable screening of expansive chemical libraries of drug-like molecules. After several iterations, we settled on a design utilizing genetic engineering to enable an HTS-compatible <u>D</u>ual <u>L</u>uciferase <u>R</u>eporter (DLR) screening assay performed in primary cells from mice expressing two distinct luminescence-based reporters **(Fig. 2A)**. Primary cells are extracted from the organ system causally linked to the disorder, such as the cortex for the *Syngap1* version of *EGS*^21, 27^. Moreover, a conditional allele was required to both enable induction of *Syngap1* haploinsufficiency and to facilitate a streamlined breeding strategy **(Fig. 2A, G)**. The first luminescence signal was envisioned as a readout of endogenous target protein through activation of the *n*BIT luciferase, a split luciferase based on an engineered variant of Nano-luciferase^28^. The second luminescence signal, generated by a firefly luciferase transgene (*f*LUC) expressed from a ubiquitous promoter, was envisioned to report global changes in protein expression and/or cellular toxicity in response to library compounds added to screening plates harboring primary cells. Inducible *Syngap1* haploinsufficiency, to enable AD disease modeling, would be achieved through a conditional *Syngap1* allele with *Lox*P sites flanking essential exons. Three distinct strains of mice were required to achieve the noted design goals, and once obtained, they would be crossed to yield offspring expressing the engineered genetic components **(Fig. 2A, G)**.

**Figure 2.**
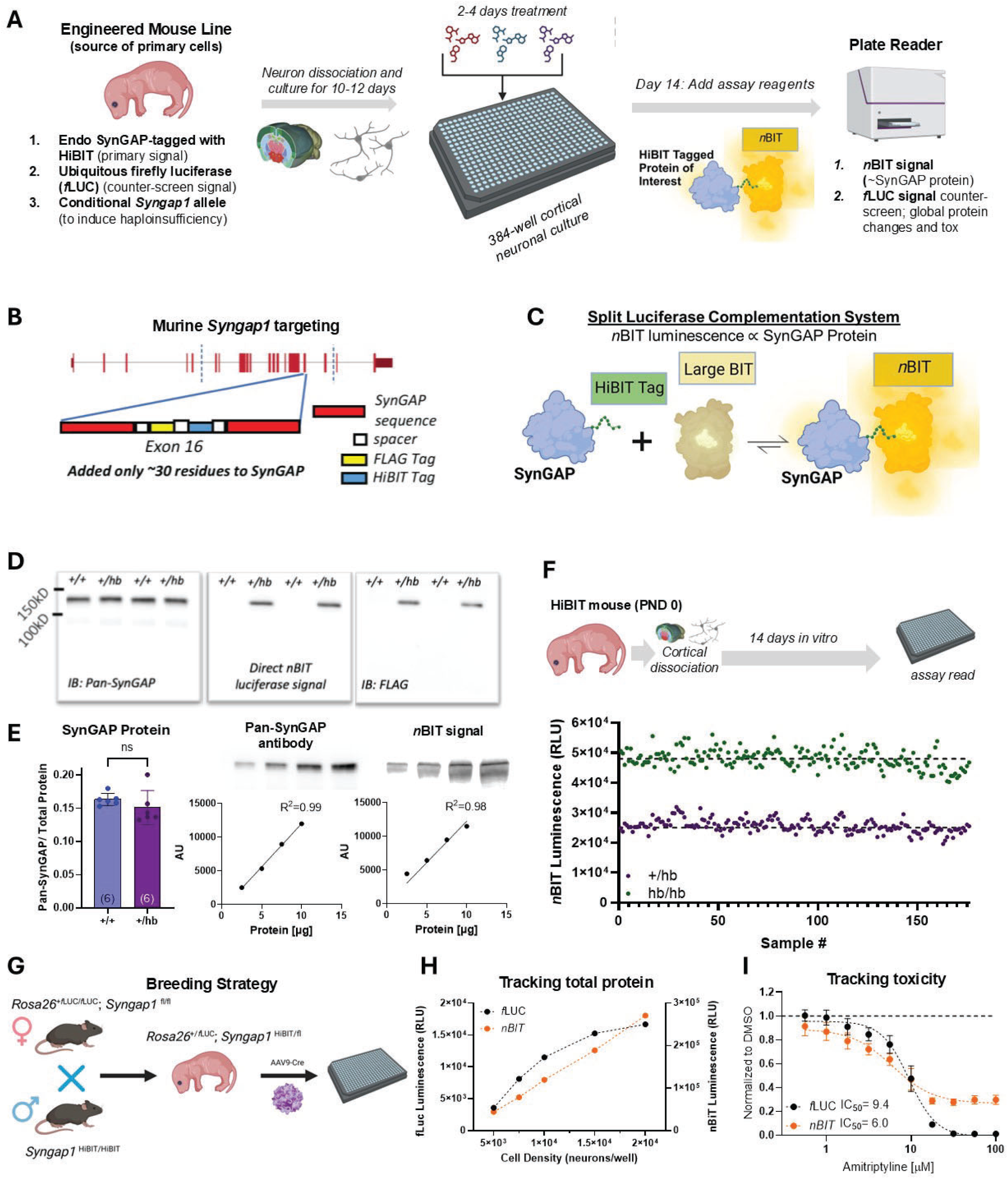
Development of primary cell <u>D</u>ual <u>L</u>uciferase <u>R</u>eporter (DLR) assay to detect changes in endogenous target protein expression. **A)** Concept of required engineered mouse line to enable endogenous SynGAP protein measurements within an HTS-like workflow. Neurons from the mouse line are cultured in 384-well plates are then treated with library compounds for 2-4 days before execution of the DLR assay. **B)** Schematic of the design and location of the HiBIT tag to be inserted into the mouse *Syngap1* gene. **C)** Endogenous SynGAP-*HiBIT* fusion protein activates *n*BIT luciferase activity. **D)** Immunoblots from PND14 mice heterozygous for the inserted HiBIT tag. Samples were probed with Pan-SynGAP and FLAG antibodies. Samples were also probed with a detection kit that enables a direct luminescent readout of proteins that contain a HiBIT tag. **E)** Quantification of endogenous SynGAP protein from immunoblots containing *Syngap1^+/+^* and *Syngap1^+/HiBIT^* mice (n=6 per genotype). Error bars represent SD. *Syngap1^+/HiBIT^* lysate was serially diluted, and probed for correlation of SynGAP levels detected using a Pan-SynGAP antibody or LgBIT. Simple linear regression was performed with Y-intercept=0, Pan-SynGAP p=<0.001, and *n*BIT p=<0.001, n=2 per concentration. F) *Syngap1^+/HiBIT^* mice were crossed to each other. *Syngap1^+/HiBIT^* and *Syngap1^HiBIT/HiBIT^* mice were cultured in 384-well plates and assayed at DIV14. **G)** Breeding strategy to yield mouse offspring to feed the *Syngap1* version of *EGS*. **H)** Neuronal plating density was varied to measure how the two DLR signals reflect changes in the amount of total protein in the well. n=128 for densities 5,000– 15,000 cells/well and n=64 for 20,000 cells/well. Error bars represent SEM. **I)** Amitriptyline, a known neurotoxic agent in primary cultured neurons, induces a dose-dependent reduction in both luciferase signals indicating that a coordinated drop in both signals reflect neuronal toxicity (n=32 per dose). Error bars represent SD.

The conditional *Syngap1* knockout mouse^29^ and a transgenic mouse ubiquitously expressing *f*LUC *(see methods)* were already constructed, validated, and available commercially. To detect endogenous SynGAP protein in mouse cells/tissues, we created a new mouse knock-in strain that expressed a *Hi*BIT ligand^11^ in-frame within protein made from the murine *Syngap1* gene **(Fig. 2B; Fig. S1A-B)**. When the *Hi*BIT tag binds to an inactive and purified fragment of Nano-luciferase, called *Large*BIT, it reactivates dormant luciferase activity **(Fig. 2C)**. The reconstituted and catalytically active *Hi*BIT-*Large*BIT complex is referred to as NanoBIT luciferase (*n*BIT from now on). Photons generated from *n*BIT activity produced from an endogenous SynGAP-HiBIT fusion protein were hypothesized to report proportional changes in endogenous SynGAP protein induced by test agents within primary cultured cortical neurons. To validate this, we first confirmed that the genetic insertion within the *Syngap1* coding region did not alter Mendelian ratios of offspring (*not shown*). We also confirmed that the insertion did not disrupt endogenous full-length SynGAP protein expression levels in mouse brain **(Fig. 2D-E)**. Insertion of the tag accurately reported relative changes in SynGAP expression within a diluted sample of cortical tissue extract. Moreover, detection of endogenous SynGAP protein through addition of LargeBIT on a traditional immunoblot predicted the levels of endogenous protein on par with antisera. We next confirmed that primary cultured neurons prepared from the *Syngap1*-HiBIT knock-in mouse enabled detection of relative changes to endogenous SynGAP protein within HTS-compatible 384-well screening plates **(Fig. 2F)**. To do this, neurons from either heterozygous or homozygous *Syngap1*-HiBIT knock-in mice were added to separate wells of the same assay plate. *n*BIT signal from homozygous knock-in neurons were on average ∼2-fold higher than signals obtained from heterozygous neurons **(Fig. 2F)**. Signals derived from *wildtype* (WT) mice (*not shown*) yielded ∼60 relative luciferase units (RLUs), resulting in a very high signal-to-noise ratio of >10e3 for detecting endogenous HiBIT-tagged SynGAP protein within primary cells or tissue. Moreover, signals derived from assay wells with either heterozygous or homozygous neurons resulted in completely non-overlapping populations **(Fig. 2F)**, indicative of low inherent assay variance. These properties suggested that this signal would be suitable to identify small molecules that regulate physiological changes in endogenous SynGAP protein within primary cortical neurons.

Primary neurons extracted from postnatal day (PND) 0 mice **(Fig. 2G)** were then cultured in assay plates to determine to what extent the non-specific *f*LUC signal reports global changes in protein expression and/or cellular toxicity. Indeed, we found that incrementally raising neuronal plating density drove a proportional increase in *f*LUC signal, confirming that this signal approximates global changes in total protein (and plating density) within assay wells **(Fig. 2H)**. This signal also reliably reported neuronal toxicity induced by amitriptyline **(Fig. 2I)**, a known neurotoxic agent^30^. As a result, the non-specific *f*LUC signal serves as a counter-screen to eliminate test agents that globally induce changes in protein expression, while also reliably reporting cellular toxicity. The former is important for rejecting test agents that fail to stimulate enrichment of the target (SynGAP) protein, while the latter is critical for directing downstream medicinal chemistry approaches aimed at improving the drug-like properties of probes that advance through a lead optimization program. Finally, we confirmed that AAV-driven Cre expression within neurons derived from this cross induced a dose-dependent decrease in SynGAP protein **(Fig. S1C)**. This confirmed our ability to induce *Syngap1* haploinsufficiency in neurons expressing the two luciferase reporters, a requirement for modeling the *Syngap1* genetic disorder in a dish.

We next optimized the assay for HTS-level scalability so that large chemical libraries of drug-like probes could be screened. Internal *in silico* modeling suggested that a strategic selection of up to 100K distinct compounds would be sufficient to sample diverse chemical space within a >2M compound collection of drug-like lead molecules available through various commercial sources *(see methods)*. To screen up to 100K compounds (in duplicate) using the primary neuron DLR assay, ∼624 assay plates would be required **(Fig. S2A-B)**. To achieve this desired level of scalability, ∼52 individual litters of mice would be needed to produce cultured cortical neurons at scale. With an average output of two batches of assay plates per week, a screen would take ∼6 months to complete. Scalability is largely dictated by the availability of mouse primary neurons. When using an optimized breeding strategy **(Fig. 2G)**, we previously demonstrated that two batches of neurons per week (*up to 32 plates*; **Fig. 1**) is possible using one full-time-equivalent laboratory position^23^. However, to minimize rates of false positive/negative data points, the assay must exhibit reliability and reproducibility, especially when performed in an iterative screening environment spanning months. Indeed, it was unclear at the time to what extent primary cortical cultures from postnatal mouse brains would exhibit the required scalability and reliability to sustain a screening platform carried out over an extended timeframe. To quantify culture reliability, plate-level quality control (QC) metrics, such as raw luciferase values from each independent signal and the coefficient of variance (CV) of these signals in negative control wells (<0.1% DMSO), were used as “pass/fail” flags for individual compound screening plates within and between culture batches **(Fig. 3A)**. In pilot studies, plates within a screening “batch” shared pooled primary neurons obtained from a single mouse litter, and as expected, QC metrics were highly similar between plates within the same batch **(Fig. 3B)**. This experiment revealed that the CV of raw luciferase values from individual plates remained within the acceptable range (<15%) even when culture density varied over a 1.5-fold range. This suggested that culture batches produced over a long timeframe within the same screening project would demonstrate the requisite reliability (e.g. few failed batches) and thus could sustain a screening platform. Consistent with this prediction, post-hoc analysis of 524 neuronal culture screening plates prepared over >1 year were found to be highly consistent, with only 4% of plates being failed (CV > 15%: **Fig. 3C)**. When plates were failed, post hoc analysis of these simple QC metrics enabled a quick diagnosis, and most “fails” were traced back to an issue with one or more channels within the robotic liquid dispenser (e.g., clogging).

**Figure 3.**
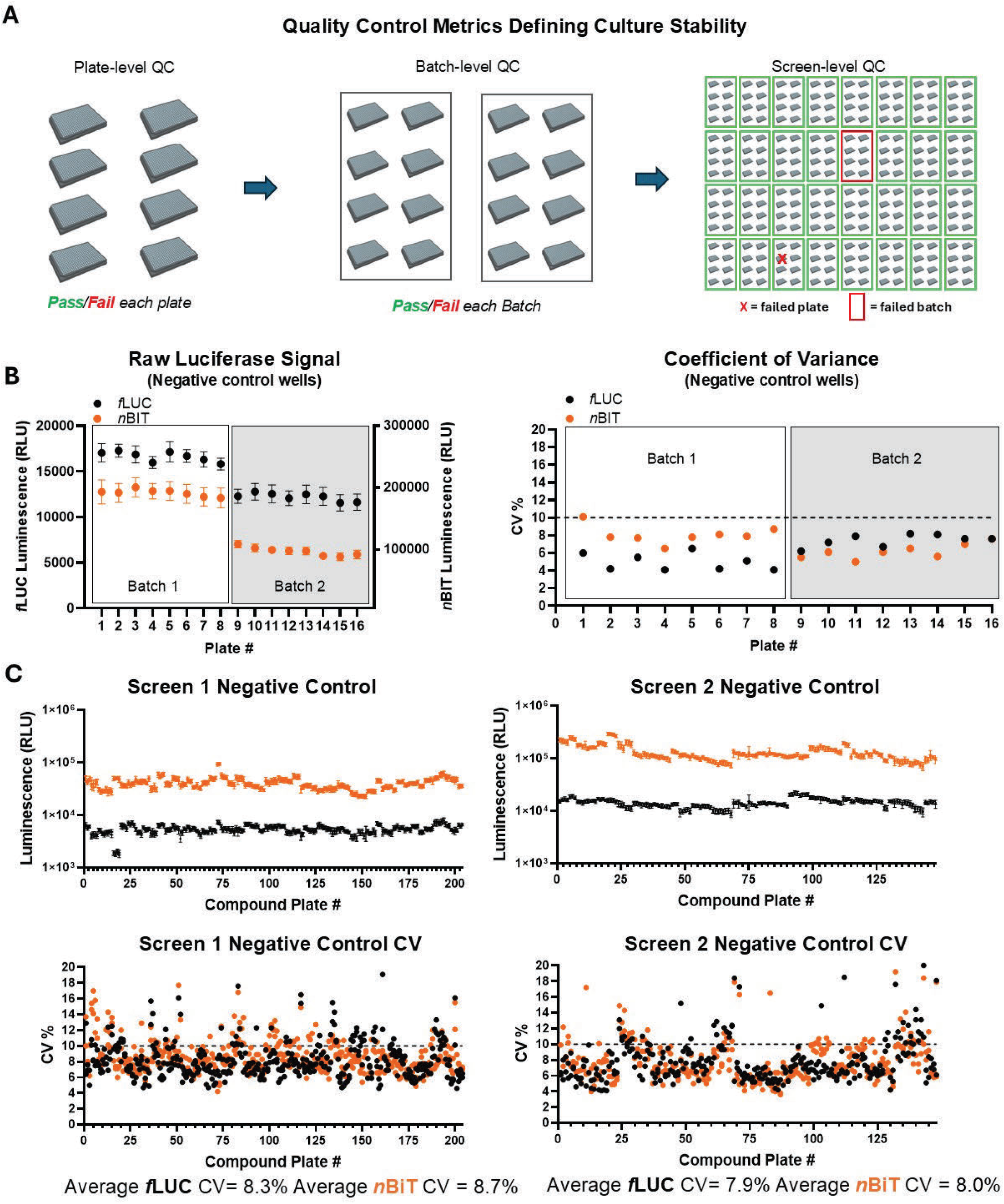
Plate- and batch-level quality control (QC) measures for screening plate data. **A)** Schematic of data hierarchy generated from the proposed iterative high-throughput screen**. B)** Luminescent values and coefficient of variance (CV) for *f*LUC (red) and *n*BIT (blue) signals across two batch of neurons plated at 15,000 cells per well (Batch 1) or 10,000 cells per well (Batch 2), n=36 per plate. Error bars represent SD. **C)** Raw luminescence values *(top)* and CV *(bottom)* for negative control wells in Screen 1 (∼59,840 compounds) and Screen 2 (∼47,360 compounds). Error bars represent SD.

### *EGS* Assays enable HTS-style screening of chemical libraries required for Lead Identification

To identify compounds that upregulate SynGAP protein expression within haploinsufficient neurons, we performed two independent screens, which together comprised >100K distinct drug-like small molecules. Two independent screens were carried out because we identified important principles in the first screen, which led to modest modifications to the screening workflow and assay parameters, which were implemented in Screen 2. For example, at the time of implementing Screen 1, there were no known small molecules that raised SynGAP protein in our culture model system. Therefore, assay plates in Screen 1 did not contain a positive control **(Fig. S2A)**. Instead, an extra row of negative controls was included. Additionally, the final compound concentration was varied so that a *post hoc* analysis of data could be used to determine how concentration impacted assay performance. In Screen 1, the concentration of library compounds ranged from ∼3.125-12.5 μM in ∼0.03125-0.125% DMSO. Moreover, compounds were pinned into primary culture assay wells and allowed to incubate for 48 hours – a notably conservative approach for inducing steady-state changes in protein abundance. The DLR assays were always carried out on day *in vitro* (DIV) 14. All compounds were screened in duplicate (N=2 screen).

To detect hits from Screen 1, an “outlier” algorithm was developed to identify compounds that selectively increased or decreased the *n*BIT signal **(Fig. 4A-C)** – the proxy measure for SynGAP protein expression in neurons. Before this could be implemented, the median *f*LUC and *n*BIT signal of the negative control wells (DMSO only) from each plate were first used as a normalization value for every individual well on each plate, including all compound wells. This enabled a z-score to be calculated, with the normalized *f*LUC signal on the X-axis and the *n*BIT signal on the Y-axis **(Fig. 4A)**. When visualized this way, both the negative control wells, as well as most compound wells, clustered near the origin. The large number of compounds interspersed with the negative controls was indicative of most compounds lacking activity in the assay, something that was expected in screens of this type, with common hit rates below 1%. However, on each plate, some wells exhibited signals that significantly deviated from the origin, which was indicative of biological activity in the assay. For example, compounds often appeared in the lower left quadrant, which was an indication of cellular toxicity. We also commonly observed a fraction of compounds that significantly increased both signals, suggesting a global increase in protein. Because the goal was to identify compounds that selectively upregulated endogenous SynGAP expression within haploinsufficient neurons, we used the normalized data to identify “Hits,” which were defined by setting 3x standard deviation (SD) threshold lines (orange vertical and horizontal lines) above and below the average of the normalized negative control wells for both *f*LUC and *n*BIT signals. Compounds showing activity above 3x SD for the *n*BIT signal and within 3x SD of *f*LUC were considered preliminary hits. We next filtered out compounds showing unacceptably high variability between replicate data points. This was done by calculating the CV of each initial compound hit, and compounds showing <10% variability across both *f*LUC and nBIT reads were retained on the final Hit List for each compound plate (**Fig. 4A** - *blue data points*). All accepted data from each compound plate data was then collapsed onto a single 2D-scatter plot **(Fig. 4B-C)**. Visualizing the complete screening data set revealed the relative position of the final Hits (*blue points*), the inactive compounds (*grey points*), and the negative controls (*green points*). Using this approach, Screen 1 yielded 127 compounds of interest across 60K compounds assayed (0.21% Hit rate).

**Figure 4.**
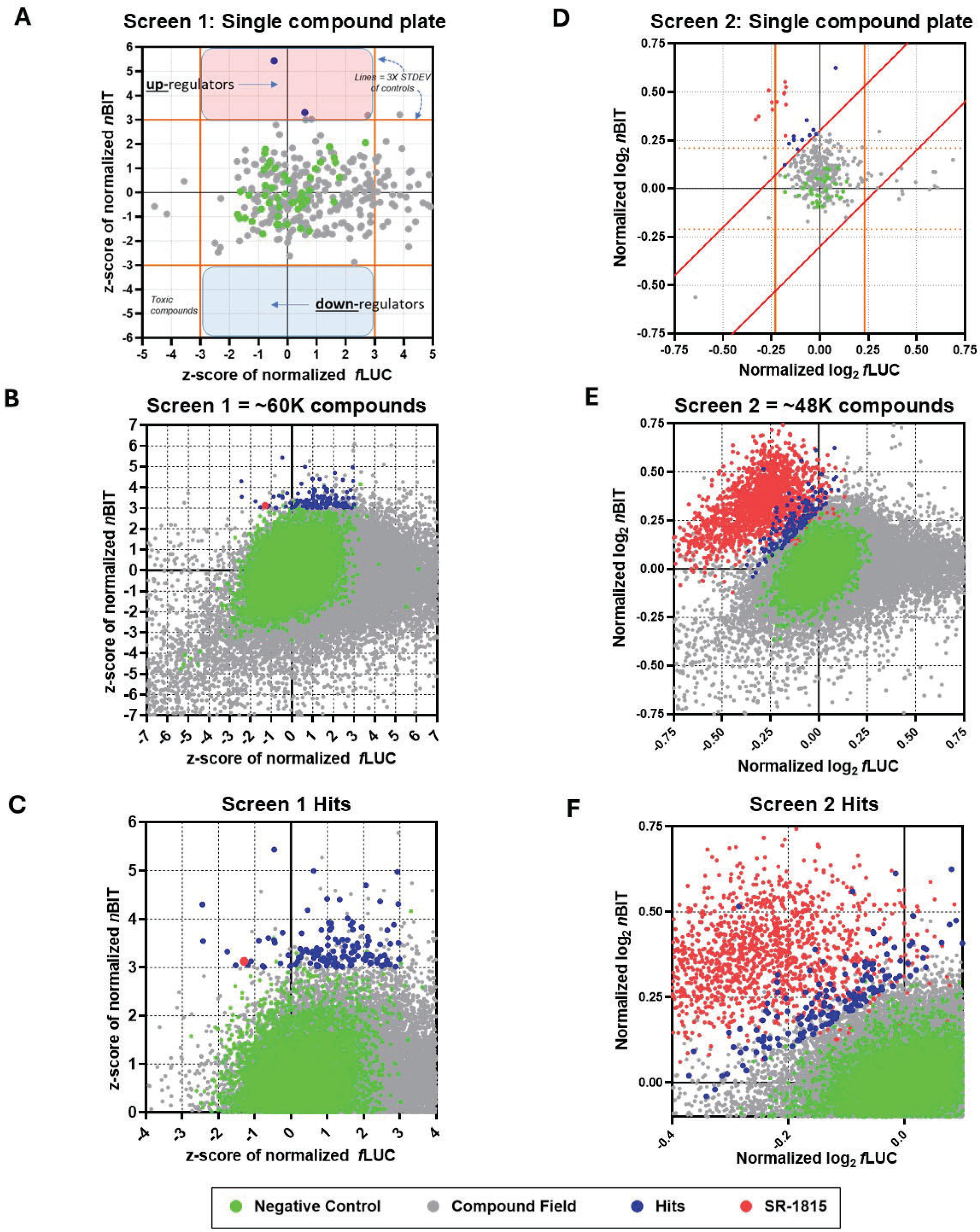
Data from two independent HTS-like screens for SynGAP protein expression “boosters”. **A)** One plate of screening data from Screen 1 visualized on a 2D scatter plot – *n*BIT (SynGAP) signal vs. *f*LUC (toxicity/selectivity) signal. Threshold lines set at 3X standard deviation of the mean for negative control wells. Red shaded section above corresponds up-regulator hit zone and blue shaded section below corresponds down-regulator hit zone. Gray dots appearing in hit zone did not meet established variability criteria. **B)** All data from Screen 1 visualized on 2D scatter plot. **C)** Zoomed in view of Fig 3B to visualize hit compounds from Screen 1 (127 compounds = 0.21% hit rate). **D)** One plate of screening data from Screen 2 with adjusted threshold lines. **E)** All data from Screen 2 visualized on 2D scatter plot. **F)** Zoomed in view of Fig 3D to visualize hit compounds from Screen 2 (171 compounds = 0.36% hit rate). For all figures, color coding refers to negative control (green), compound field (gray), hits (blue), and SR-1815 (red).

We next screened a distinct library of ∼48,000 compounds that were pre-selected based on chemical features known to be conducive to developing neuroactive small molecule therapeutics. For this second screen (e.g., *Screen 2* – **Fig. 4D-F**), several workflow improvements were implemented, which were inspired by experience gained from Screen 1. Consequential modifications included a single screening concentration (∼3.125 μM), doubling the compound incubation time to 96hrs (DIV10-14), and incorporation of a positive control on all screening plates **(Fig. S2A)**. The positive control was discovered during Screen 1 **(Fig. 4B** – *red dot*). Including this positive control on screening plates led to an improved computational approach for identifying compounds with significant activity in the assay **(Fig. S3A-D)**. By quantifying the pattern/spread of positive control, negative control, and inactive compound data points across all screening plates, a clear positive correlation between *f*LUC and *n*BIT signals in each of these three populations was identified. This relationship functioned as a type of Loading Control for each well, which allowed us to estimate the relative difference in the amount of neuronal material from well-to-well, and was instrumental in developing an improved hit detection algorithm (*denoted by sloped lines;* **Fig. 4D)**. The preliminary Hits were next filtered for obvious false positives by assigning a Non-Repeatedness value. Non-Repeatedness represents a single numerical value that considers both the embedded positive relationship between *f*LUC and *n*BIT signals and the continuity of the duplicate data points afforded by the N=2 screen design **(Fig. S3E-F).** Implementation of the new detection algorithm and filtering for Non-Repeatedness revealed 171 additional Hits in Screen 2 (0.36% hit rate) (*blue dots;* **Fig. 4E-F).** Several Hits met, and in some cases exceeded, the performance of the positive control. This, combined with a similar hit rate, suggested that the somewhat narrower chemical space within CNS-focused library did not negatively impact overall screening performance, which may be explained by the improved assay workflows implemented in Screen 2.

### *EGS* assays support biological validation of preliminary screening Hits

An essential component of the *EGS* platform is a workflow dedicated to validating the most promising compounds of interest identified in HTS-style screens **(Fig. 1**; **Fig. 5A)**. The goal of this workflow is to develop extreme confidence that a preliminary Hit obtained in the original screen upregulates endogenous steady-state SynGAP protein abundance within *Syngap1* haploinsufficient cortical neurons. Of the several validated probes at the time of submission, SR-1815 **(Fig. 3B-C**; *red dot*) was chosen as a proof-of-concept molecule to demonstrate the effectiveness of the Hit Validation Workflow. Indeed, searches of the literature (Scifinder, Pubmed) were conducted for SR-1815 analogs and identified no known chemistry, biology, or pharmacology related to this scaffold. The compounds were not represented in the public domain other than as analogs in a screening collection. Thus, if validated, SR-1815 would become a first-in-class drug-like small molecule that upregulates SynGAP protein in a disease-modeling context. The probe validation workflow was comprised of up to five additional levels of validation beyond the initial screening data **(Fig. 5A)**. The first steps in the validation pipeline were designed to determine to what extent a compound regulates the *n*BIT signal in a dose-dependent manner. SR-1815 demonstrated dose-response activity from both the serial dilution of the original library material *(not shown)* and from freshly sourced compound “powder” **(Fig. 5B)**. To rule out a direct effect of the small molecule on *n*BIT enzymatic activity, we utilized a cell-free counter-screen, where LargeBIT, a purified Halo-HiBIT fusion protein, and the compound of interest were added together with the assay reagents. In this cell-free assay, SR-1815 did not increase *n*BIT activity at any concentration tested **(Fig. 5C),** suggesting that it may regulate SynGAP protein expression.

**Figure 5.**
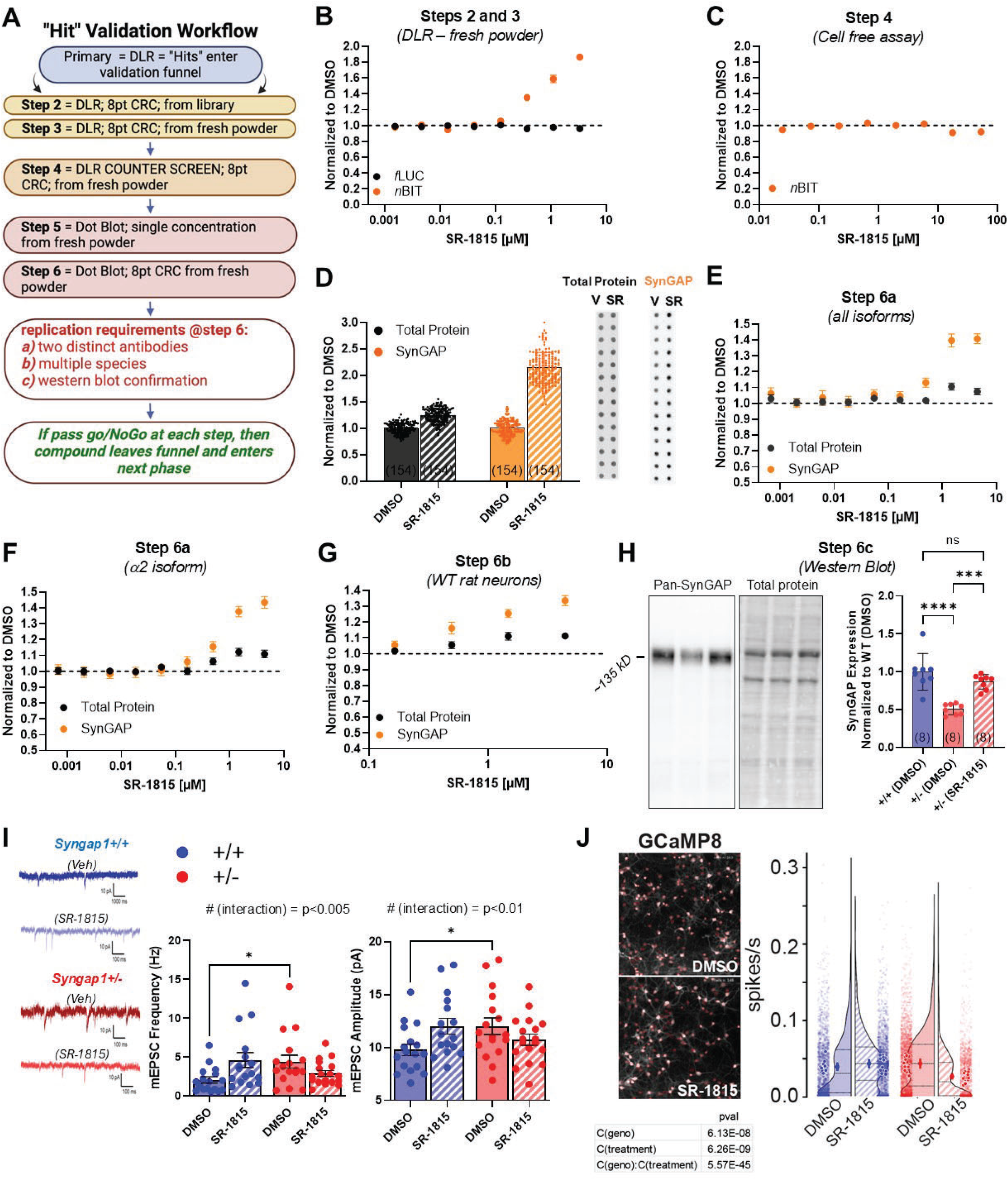
Workflow for validating preliminary hits identified in *EGS* HTS-style screen. **A)** Validation workflow encompassing luciferase-based assays, counter-screens, and protein validation assays. **B-C)** SR-1815 in DLR or *n*BIT counterscreen 8-point dose-response assay. n=12 per dose for DLR and n=10 per dose for cell free assay. Error bars represent SEM. **D)** Single dose Dot Blot experiment with SR-1815 treated at 1.5 µM for 14 days yielding signals for total protein and SynGAP protein expression A Mann-Whitney test was performed for total protein: U=1609, p=<1.0*^e^*15, and SynGAP U=0, p=<1.0*^e^*15. Error bars represent SEM. **E-F)** Dot blot dose response of SR-1815 treated for 7 days utilizing a SynGAP antibody that recognizes either all isoforms (E) or only the α2 C-terminal isoform (F); n=28 per dose. Error bars represent SEM. **G)** Dot Blot experiment for SynGAP protein expression in WT rat neurons. Seven-day treatment (DIV 7-14); Pan-SynGAP antibody; n=28 per dose. Error bars represent SEM. **H)** Western blot with samples extracted from primary cultures derived from either *Syngap1^+/+^* and *Syngap1^+/-^*. Neurons were treated for 14 days with either DMSO or SR-1815 (1.5 μM); One-way ANOVA with Tukey’s post hoc tests were used. F(2,21)21.46, p=0.000008, n=8 per treatment. Error bars represent SD. **I)** Representative traces from mEPSC recordings in *Syngap1^+/+^* and *Syngap1^+/-^* neurons treated with vehicle or SR-1815 *(left)*. Plot showing mEPSC frequency for the four different conditions (*middle*; 2-way ANOVA with Fisher’s LSD post hoc test, Genotype: F (1,64) = 0.2557, p = 0.6148, Treatment: F (1, 64) = 0.6138, p = 0.4362, Interaction: F (1,64) = 9.168, p = 0.0035). Plot showing mEPSC amplitudes for the four different conditions (right; 2-way ANOVA with Fisher’s LSD post hoc test, Genotype: F (1, 64) = 0.5606, p = 0.4568, Treatment: F (1, 64) = 0.5601, p = 0.4570, Interaction: F (1, 64) = 7.344, p = 0.0086). n= 17 for *Syngap1^+/+^: vehicle;* n= 16 for *Syngap1^+/+^: SR-1815;* n= 17 for *Syngap1^+/-^*: vehicle; n= 18 for *Syngap1^+/-^* : SR-1815. Error bars represent SEM. **J)** *Syngap1+/+* and *Syngap1+/-* neurons transduced with AAV9 vectors expressing Flex-gCAMP8f and Cre (to control labeling). Neurons were treated with DMSO or 1.5 μM SR-1815 for 14 days. Calcium imaging performed on DIV14. Plot showing the spiking frequency (spikes per second) for the four different conditions (OLS regression with Tukey HSD post hoc test, Genotype: F(1,6792)=29.4, p=6.1e-8, Treatment: F(1,6792)=33.8, p=6.3e-9, Interaction: F(1,6792)=200.9, p=5.5e-45). For DMSO control 12 fields were imaged for *Syngap1^+/+^ Syngap1^+/-^* and for SR-1815 respectively 14 and 17 fields for *Syngap1^+/+^ Syngap1^+/-^* were imaged from at least 4 wells per conditions. The total number of segmented neurons were DMSO *Syngap1^+/+^*: 1693, *Syngap1^+/-^*: 2047, SR-1815 *Syngap1^+/+^*: 1803 and *Syngap1^+/-^:* 1253.

Given these results, we hypothesized that the compound stimulates SynGAP steady-state protein expression in neurons. Evidence supporting this hypothesis necessitated the development of a scalable orthogonal assay that directly measures SynGAP protein. A scalable Dot Blot protein assay was developed that combined a 384-array pin tool, a nitrocellulose membrane, a label that reports total protein, and a series of knock-out validated SynGAP antibodies **(Fig. S4A)**. After neuronal lysis directly within the 384-well plate, samples were incubated to fluorescently label lysine residues as a measure of total protein using derivatizer, 3-(2-Furoyl)quinoline-2-Carboxaldehyde, and activator, Mandelonitrile. Lysate from each well was pinned onto a nitrocellulose membrane. After optimization, this technique reliably reported accurate levels of total protein in each sample on the membrane **(Fig. S4A)**. The membrane was also exposed to a monoclonal antibody that detects a motif expressed in all SynGAP protein isoforms (e.g., pan-SynGAP). After washing and secondary antibody exposure, the membrane was imaged for total protein and incubated with chemiluminescent substrate. This resulted in a strong signal that was linear across protein concentrations that spanned an order of magnitude **(Fig. S4A)**. The Dot Blot assay was then pressure tested to determine how well it measures changes in steady-state SynGAP protein levels in primary cortical neurons. To do this, we utilized an already validated *Syngap1* conditional rescue mouse line^29, 31^, which was engineered to express an artificial exon containing a stop codon and an artificial poly-A sequence within the mouse *Syngap1* gene. This exon is efficiently spliced into *Syngap1* transcripts, which prevents SynGAP protein expression. However, this exon is flanked by LoxP sites, and therefore expression of Cre recombinase re-activates SynGAP protein expression due to excision of the artificial exon **(Fig. S4B)**. It was previously shown that mice heterozygous for this targeted allele have roughly half SynGAP protein expressed in neurons relative to WT littermates, while homozygous mice nominally express SynGAP protein^29^. This Dot Blot technique accurately reported the known expression changes of SynGAP in neurons derived from this mouse line, with nominal expression of SynGAP protein in homozygous neurons and roughly half the normal levels of protein expressed in the heterozygous neuronal population **(Fig. S4C)**. Importantly, the Dot Blot was also able to detect the Cre-dependent re-expression of SynGAP protein in both homozygous and heterozygous neurons **(Fig. S4C)**. Furthermore, the Dot Blot assay was further validated using antisera that detects three of the four major SynGAP C-terminal isoforms **(Fig. S4D)**. This is consequential because these isoforms have unique spatial/temporal expression profiles and distinct biological functions *in vivo*^32,33^. Finally, we developed custom software that automated analysis of Dot Blots **(Fig. S5A-D; Fig. S6A-B)**, which dramatically increased the scalability of SynGAP protein detection.

Using the now-validated Dot Blot assay, we found that SR-1815 could double endogenous SynGAP in heterozygous neurons compared to DMSO controls **(Fig. 5D).** Moreover, the compound stimulated a dose-dependent increase in SynGAP protein as measured by both a pan-SynGAP antibody that detects all isoforms **(Fig. 5E)**, and an antibody that recognizes only the α2 isoform **(Fig. 5F)**. Given that the anti-α2 signal matched the anti-pan-SynGAP signal within the same samples, this result strongly indicated that SR-1815 stimulates relatively equal expression of all SynGAP C-terminal isoforms^32^. Pairing optimized assay conditions with a traditional Western blot technique, SR-1815 rescued SynGAP protein levels in heterozygous KO neurons **(Fig. 5H)**. The compound also stimulated SynGAP expression in neurons derived from typically developing *WT* rats **(Fig. 5G)**. This demonstrated that SR-1815 efficacy is not limited to mouse neurons and the compound is effective in both a typically developing genetic background and in a background of *Syngap1* haploinsufficiency. Collectively, these data demonstrate that *EGS* validation workflows can identify compounds from HTS-style screens that reliably stimulate protein expression driven by the pre-selected genetic target.

Given that SR-1815 approximates rescue of SynGAP expression in *Syngap1* haploinsufficient cultured cortical neurons, we next evaluated to what extent the compound impacted consequential phenotypes caused by *Syngap1* haploinsufficiency. SynGAP heterozygosity has been shown to increase the frequency and amplitude of miniature excitatory postsynaptic events (*m*EPSCs) in primary cultures^22, 34^, which is a measure of excitatory synapse strength. Therefore, we designed an experiment that enabled an assessment of how excitatory synapse strength was impacted by SR-1815 in both *WT* and *Syngap1* haploinsufficient neurons. We observed a strong interaction between drug and genotype for both *m*EPSC frequency (p<0.005) and amplitude (p<0.01), demonstrating that these effects of the compound were genotype-dependent **(Fig. 5I)**. Post-hoc comparisons revealed that both *m*EPSC frequency and amplitude were significantly increased in neurons derived from *Syngap1* heterozygous animals, a result that agreed with past studies. The strong interaction was driven by the opposing effect of the compound on mEPSC measures within each genotype. The compound tended to increase excitatory synapse strength in *WT* neurons but tended to decrease it in haploinsufficient neurons. These opposing trends are what drove the strong interaction in the statistical model. Thus, while the effect of the treatment fell short of *post hoc* significance within each genotype, the strength of the interaction overall provides strong evidence that the compound has genotype-specific bidirectional effects on excitatory synapse function (e.g., increases synapse strength in WT neurons; decreases synapse strength in haploinsufficiency neurons). This is important because *SYNGAP1/Syngap1* haploinsufficiency causes neural hyperexcitability and seizures in humans and rodents^21^. Moreover, reducing excitatory synapse strength through AMPA receptor inhibition improves cognition-linked brain rhythms in *Syngap1* heterozygous mice^35^.

The observed compound-induced genotype-specific effects on excitatory synapse strength suggested that SR-1815 may also regulate neuronal activity in a genotype-specific manner. To test this, GCaMP8 dynamics were measured across thousands of individual neurons treated with vehicle or SR-1815 from cortical cultures derived from each genotype **(Fig. 5J)**. Overall, the effect of SR-1815 within each genotype was consistent with results obtained from measurements of excitatory synapse strength. For example, this analysis detected a main effect of genotype (p<0.001), and a posthoc comparison confirmed that vehicle-treated heterozygous neurons had increased activity compared to vehicle-treated WT neurons (p<0.001). Moreover, there was a main effect of treatment (p<0.001), indicating that the SR-1815 significantly regulated spike rates in neurons from both genotypes. However, a significant interaction was detected between genotype and treatment (p<0.001), demonstrating that the compound regulated activity in a genotype-specific manner. Indeed, SR-1815 significantly *<u>increased</u>* activity in *WT* neurons (WT-DMSO vs. WT-drug, p<0.001), while it substantially *<u>decreased</u>*it in heterozygous neurons (Het-DMSO vs. Het-drug, p<0.001). Thus, the compound drove normally hyperactive heterozygous neurons to activity levels at or below that of *WT* neurons. Given the potential significance of this result, we repeated this experiment in heterozygous neurons, but this time using multiple doses. Importantly, a dose-dependent decrease in neuronal activity by SR-1815 was observed in this additional experiment **(Fig. S7),** demonstrating that adjusting the dose of the compound can tune hyperactive heterozygous neurons to levels approximating the *WT* state. Taken together, *EGS* can identify a drug-like probe that raises endogenous expression of the targeted AD gene, and this probe can counteract the functional consequences of genetic haploinsufficiency within a disease modeling cellular context. Moreover, the genotype-specific effects of SR-1815 highlight the importance of working in the appropriate cellular contexts when exploring the function of phenotypic probes.

### *EGS* assays facilitate preclinical drug development

*EGS* yields drug-like probes that intersect with AutD gene biology and function. Therefore, it is critical to demonstrate that the platform can support preclinical drug development workflows. An initial first step in the drug development pipeline is to optimize the lead candidate scaffold through synthetic chemistry. As a first step in this process, medicinal chemists often perform a “shotgun” approach, where substitutions are made throughout the different motifs within the compound to identify the positions on the molecule that can tolerate significant modifications.

We developed an optimized workflow that can jumpstart preclinical development by bypassing this initial random “shotgun” approach to scaffold modification. Instead, our approach is to first identify and then purchase small quantities of close analogues of a lead molecule that exist within the extensive small molecule collections held by various commercial partners. DataWarrior^36^ is an open-source cheminformatics tool that can interface with the full compound collection of various commercial entities that source screening collections for drug discovery. This tool is used to identify compounds within the full collection that are chemically similar to a lead originally identified from the screening library **(Fig. 6A)**. To begin the SR-1815 development process, we identified 200 structurally-related compounds within the full collection. Next, these compounds were sorted based on structural diversity. This ensures that none of the structural motifs within the lead compound are overweighted in the identification process. Then, 100 compounds that exhibited the broadest structural diversity across the pool of 200 selected compounds were sourced. This process yielded 100 SR-1815 analogues that resembled first-order derivatives of the lead **(Fig. 6B)**, many of which would have been made by a medicinal chemist in a traditional “shotgun” approach. The selected compounds were sourced from the vendor, usually delivered within three weeks, and then tested for dose-response activity in the *Syngap1* DLR assay **(Fig. 6C)**.

**Figure 6:**
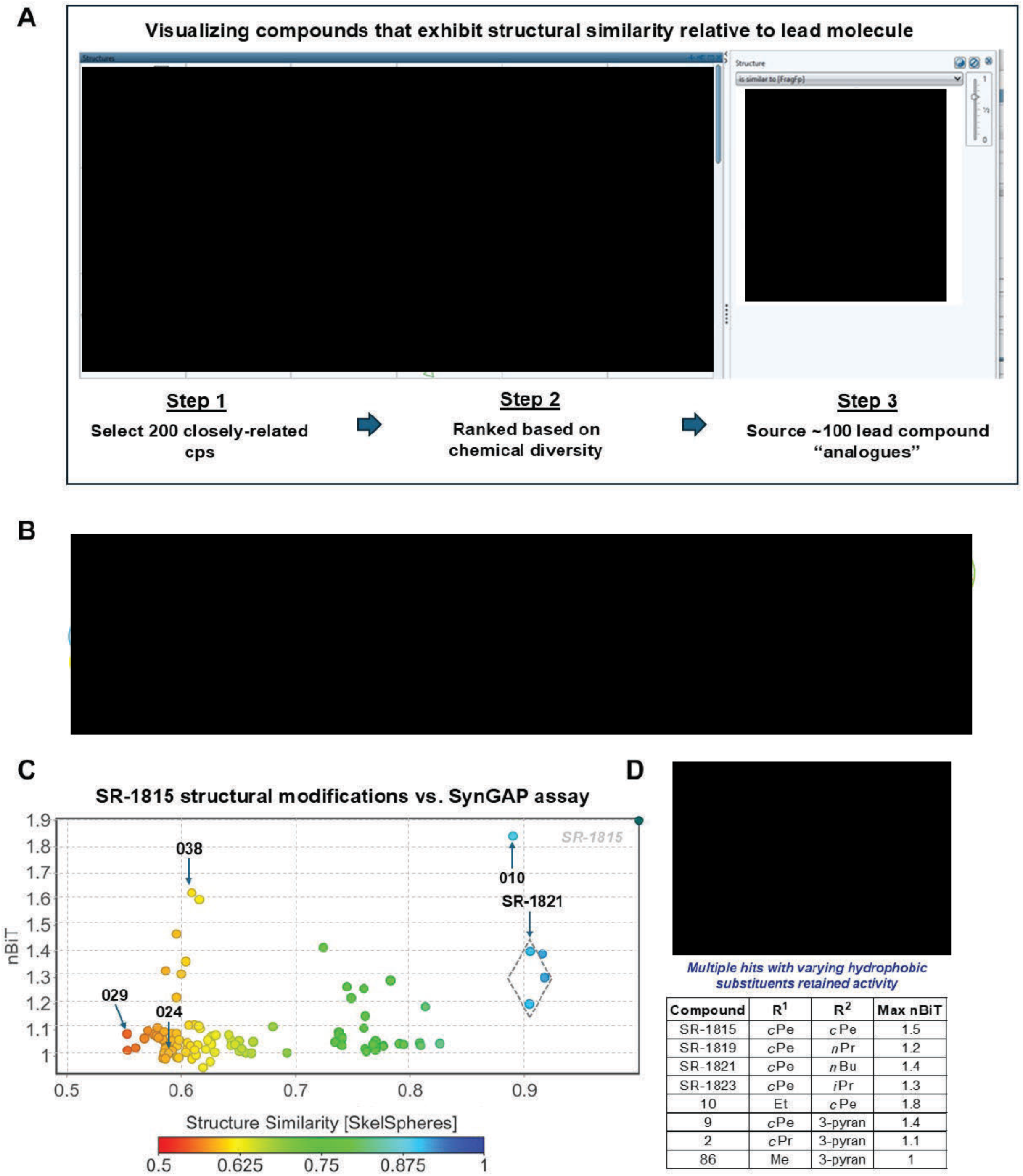
Accelerating Preclinical Development of Lead Compounds Identified through *EGS*. **A)** *Top,* Screen capture from DataWarrior exploration of SR-1815-like compounds within the full collection of small molecules. *Bottom*, Process for selecting 100 structurally similar SR- 1815-like compounds. **B)** Visualization of relative structural diversity among identified SR-1815- like compounds. Colored circles reflect different core motifs within SR-1815. **C)** Relationship between molecular structure and SynGAP assay activity from 104 SR-1815-related compounds. The 104 compounds are comprised of 100 close analogues of SR-1815 (panel A), 3 related compounds in the original screening library (e.g, SR-1821; grey circle), and SR-1815 (reference compound). **D)** Identification of clearly modifiable R-groups derived from structure-activity analysis of SR-1815 analogues shown in panel (C).

Three of the compounds flagged in the cheminformatic profiling were present in the original screening library – SR-1819, SR-1821, SR-1823 – and each analogue retained activity in the SynGAP DLR assay. One of these compounds, SR-1821, was flagged as a hit in the original screen, which supports the validity of this profiling approach. Importantly, analogues that retained activity were further validated in the Dot Blot protein assay (*not shown*). Moreover, the retained activity from these three analogues indicated that the 5-position of the pyrazole (R^2^) was amenable to substitution **(Fig. 6D)**, allowing for cyclic alkanes and ethers as well as linear and branched alkanes. Analyzing DLR data from the other 100 related compounds revealed additional areas of the compound that could sustain substitutions **(Fig. 6B-C)**. The pyrazole ring was difficult to replace with many group substitutions rendering the molecule completely inactive. However, some substituted imidazole derivatives did retain activity, indicating that a free NH group was not required. 1-N-alkyl substitution of the urea was tolerated with a range of groups from ethyl and cyclopropyl to cyclopentyl affording active molecules (R^1^). It is not yet clear if the 3-NH-urea is required for activity or the cyclic urea itself because the molecules available for purchase to test these positions were not available and will need to be synthesized. The C5 amide appears to be important for activity as well, but modifications including the reverse amide, amines, ethers and cyclic versions were not available for purchase. This strategy will be addressed during traditional SAR studies. In summary, the “SAR by purchase” approach greatly facilitated the development plan for how and where substitutions can be made to potentially optimize potency and efficacy within identified phenotypic probes. The ability to identify and then quickly receive 100 analogues of the original library molecule greatly accelerated the SAR program and validated SR-1815 as a viable preclinical lead candidate. Finally, this experiment demonstrated that *EGS* assays are sufficiently scalable and reliable to direct an SAR preclinical development program.

## DISCUSSION

*EGS* is a significant technological advance because it provides a platform to identify drug-like small molecules that regulate endogenous expression of proteins that directly regulate cellular states, such as the shift from health to disease **(Fig. 7A)**. Here, we show through HTS-style phenotypic screening, that *EGS* yielded many small molecules suggestive of raising SynGAP expression, including the extensively validated lead molecule, SR-1815. SR-1815 was the first molecule that we attempted to fully validate. Its ability to raise SynGAP protein in a relevant cellular context, combined with its ability to mitigate disease-associated functional phenotypes in *Syngap1* haploinsufficient neurons, is strong evidence that the EGS platform yields useful phenotypic probes. Moreover, EGS assays were essential during molecular deconvolution and mode of action studies of SR-1815 (**Fig. 7B**; *see companion paper Douglas et al.,* BioRxiv, *2025*). In addition, we show that EGS assays also promote preclinical development of phenotypic probes discovered using this platform. Together, these successes indicate that EGS is a powerful platform for discovery of phenotypic probes that regulate expression of proteins that cause cellular disease states. In support of this, several additional compounds of interest that were identified in the primary screens have also been validated. These compounds are under development through parallel research programs. Finally, nearly 200 Hits have yet to undergo validation, suggesting that additional SynGAP upregulating probes will be discovered in the near future.

**Figure 7:**
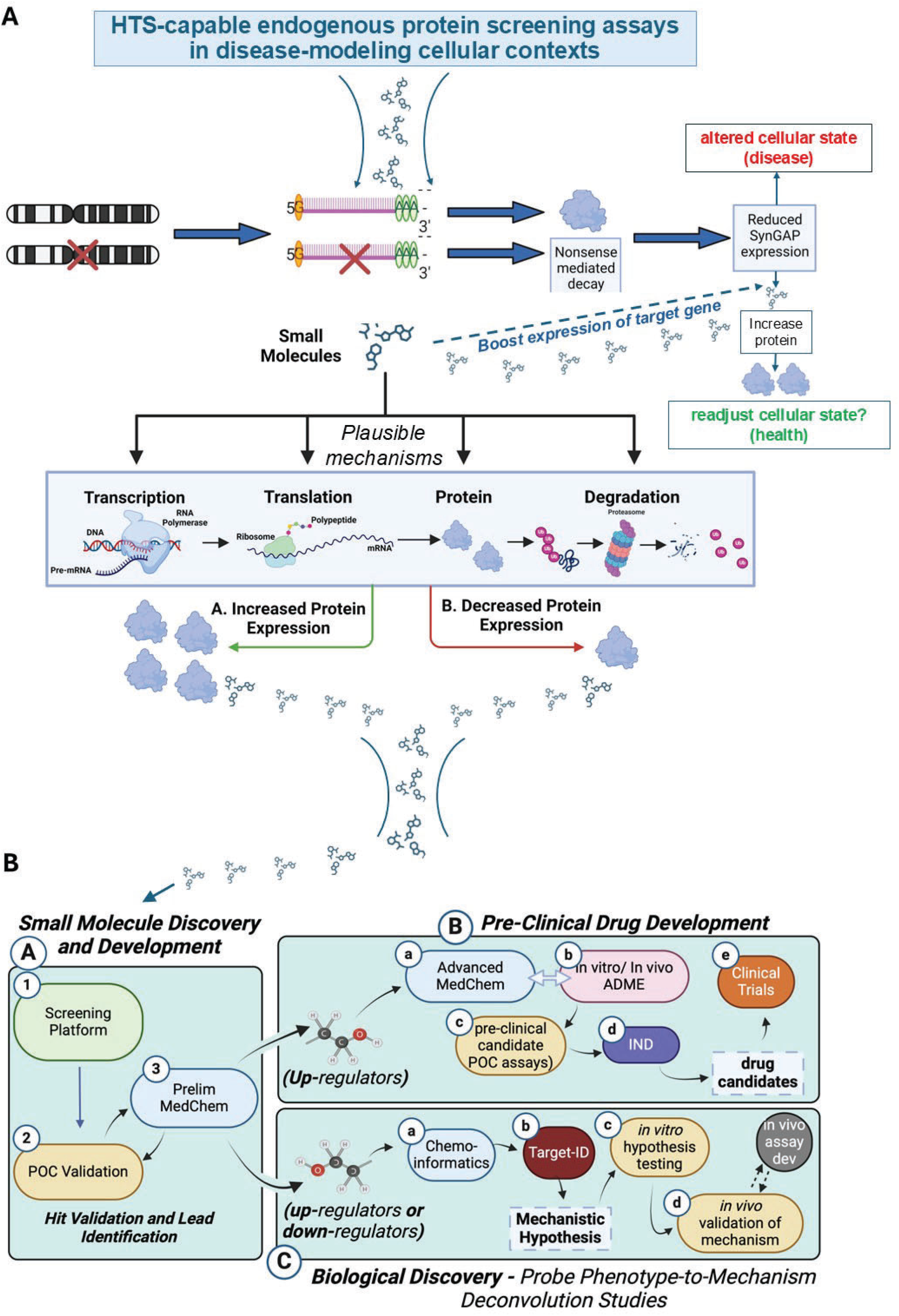
Complete *EGS* Workflow. **A)** Entry point for platform is to choose a gene and then screen chemical or biological agents using HTS-compatible assays that read-out endogenous protein expression of the gene of interest within disease-modeling state. The test agents could increase protein expression in a variety of ways, with the expected outcome of boosting protein resulting in a switch from “disease” to “healthy” cellular state. **B)** (*a, left box*) Molecules suitable for further study enter the development pipeline workflows. A compound travels through two distinct sub-paths: (*b*, *top right box*) the probe undergoes pre-clinical development toward eventual IND application; (*c, bottom right box*) probe enters phenotype-to-mechanism studies aimed at identifying its molecular targets and how it regulates signaling to switch the state from “disease” and toward “health”.

SR-1815 is a pivotal discovery; no small molecule had previously been identified as capable of restoring SynGAP levels in a haploinsufficient state. SR-1815 represents a first-in-class proof-of-concept molecule. Moreover, SR-1815 is the first small molecule capable of counteracting both the root cause of *Syngap1* haploinsufficiency (e.g., low neuronal protein expression), and a critical functional consequence of it (neuronal hyperexcitability). The probes discovered through this platform offer unique translational opportunities. The current state of the art in treating haploinsufficiency disorders is to develop targeted ASOs that disinhibit mRNA-dependent gene suppression mechanisms or to utilize viral vectors that either edit or replace a dysfunctional gene copy^37^. While these approaches have shown successes for some genetic disorders, they do not always successfully translate, nor are they necessarily the best treatment modality for every patient. For example, ASOs are not brain penetrant and must be given intrathecally to treat CNS disorders, which introduces safety concerns and discomfort for the patient^38^. Further, ASOs have half-lives of weeks and viral-based clinical approaches lack effective shut-off mechanisms, resulting in challenges related to tuning the effectiveness of these therapeutic approaches. It is well established that genes that cause AutD disease require tightly controlled expression levels to maintain cellular health^39^. Too little or too much expression of these powerful genes can, on their own, cause disease states. Small molecules remain the gold standard therapeutic agent for treating neurological disorders because they can be improved through chemistry to achieve brain penetrance and optimized delivery route (oral, IV). In addition, their levels in the body can be easily adjusted through dose titration. In certain situations, small molecules can be viewed as a superior first-line initial treatment option for genetic disorders^40^. At the very least, small molecule development provides an additional strategy for treating genetic loss-of-function disorders, which can greatly benefit patient populations.

The power of phenotypic screening lies in the ability to discover probes that drive disease-associated phenotypes of interest through complex, multi-target mechanisms. However, the major drawback to phenotypic screening is the lack of target-based information that mechanistically link the discovered probe to the phenotype of interest. EGS was conceptualized from the outset with the understanding that discovering the mode of action of newly discovered probes would be fundamental to the success of the platform **(Fig. 7B)**. Our companion paper, Douglas *et al. 2025* (Biorxiv), combined *EGS* assays with molecular-genetic and novel chemo-proteomics approaches, such as MesoMap, to create a workflow that successfully deconvolved SR-1815’s molecular targets. We found that it is a multi-kinase inhibitor, and inhibition at multiple kinase targets appears to contribute to its ability to raise neuronal SynGAP expression. The kinase inhibition profile of SR-1815 was related to several FDA approved anticancer medicines. This led to two additional discoveries. First, SR-1815 exhibited anti-cancer activity. Second, FDA-approved anticancer therapeutics also raised SynGAP expression. Thus, repositioning anticancer therapeutics may accelerate treatments for *Syngap1* genetic disorders, while the SR-1815 series could be additionally developed into a novel cancer therapeutic. These findings emphasize the power of phenotypic screening approaches in general, and how focusing on target deconvolution can broaden the potential therapeutic scope of probes discovered using EGS.

Selection of the cellular context is an important factor related to the success of phenotypic screening. In the *Syngap1* version of EGS, we chose mouse primary cortical neurons from *Syngap1* haploinsufficient mice over patient-derived human neurons as the screening context. This was done for several reasons. First, primary cortical neurons have been, and continue to be, the gold standard *in vitro* discovery model for understanding neuronal cell biology, especially the processes related to synapse biology and how synapse biology regulates network activity^43, 44^. Second, there is overwhelming evidence supporting the fundamental function of SynGAP expression in cortex and in cortical neurons. *SYNGAP1*-DEE patients express irregular cortical EEG rhythms^17, 18^ and altered cortical sensory processing^45^, which when combined with reports of disrupted higher cognitive functions^14^, support the role of this protein in regulation of cortical neuron function. *Syngap1* haploinsufficient mice and rats model domains of brain dysfunction observed in human *SYNGAP1* patients^46, 47^. In mouse models, regulation of *Syngap1* expression selectively within cortical neurons is both necessary and sufficient to modulate disease-associated phenotypes^21, 27^, including higher cognitive functions and seizure susceptibility caused by neural hyperexcitability. Cultured primary cortical neurons derived from *Syngap1* haploinsufficient mice express analogous phenotypes, including synaptic^22,34^ **(Fig. 5I)** and cellular^48^ **(Fig. 5J)** hyperfunction. Indeed, these key phenotypes were ameliorated by SR-1815. Third, human excitatory neurons derived from patient iPSCs are substandard for replicating disease-linked biology related to impaired function of synapse dynamics, neural plasticity, and the emergent phenotypes that arise from these complicated processes^43^. Fourth, the scalability of iPSC-induced neurons is severely limited relative to mouse primary neurons because the former develop much more slowly. It can take at least one month to achieve reliable synapse function in induced neurons, and to achieve functional synapses, they must be co-cultured with astroglia, which further impacts economies of scale. For certain phenotypes, iPSC-derived neurons may be the more appropriate choice, such as regulation of neurite outgrowth^49^. However, for complicated phenotypes that integrate cellular processes linking synapse function and plasticity with network dynamics, primary mouse neurons can often be the superior cellular context.

EGS is not limited to identification of small molecule “boosters” linked to genetic loss-of-function disorders. Rather, EGS was developed to be modular and flexible, where the FLAG-HiBIT tag can be inserted into any gene of interest using a straightforward gene editing approach. When using mice as a model system, development of the new HiBIT tagged mouse line can be bred to the existing *f*LUC-expressing line, leading to an in-mouse DLR assay that targets any gene of interest. Similar strategies could be applied to endogenous proteins expressed in induced cells derived from patient iPSCs. As a result, DLR assays within models characterized by either low or high/toxic proteins can be developed to carryout endogenous screening within relevant cellular contexts. This can be particularly useful for neuropsychiatric disorders. For example, the MECP2 gene causes Rett Syndrome when it is expressed at low levels, yet causes a distinct brain disorder when this same gene is overexpressed through genetic duplication, and excellent models exist for each disorder^41, 42^. A single EGS platform screen for endogenous regulators of MECP2 protein expression in *WT* cortical neurons would, in theory, identify both up- and down-regulating compounds that could be tested in models for both disorders. In support of this, while not the focus of our screen, we did identify several candidate SynGAP down-regulators and validated at least one of them using our EGS approaches (*not shown*).

## METHODS

### Mice

All mouse procedures were conducted in accordance with the NIH Guide for the Care and Use of Laboratory Animals, and all methods were authorized by the Scripps/UF Scripps Biomedical Research Institutional Animal Care and Use Committee. Both males and females (M/F) were used in all experiments. The design and maintenance of the constitutive *Syngap1* KO and two conditional *Syngap1* lines have been described previously ^29^ and are available at Jackson Labs (KO=#008890; cKO =#029303; cRescue=#029304). The firefly luciferase mouse is also available from Jackson Labs (008450). *Syngap1*-HiBIT knock-in strain was created by standard CRISPR methods in collaboration with the Salk Institute Transgenesis Core facility. Briefly, mouse blastocysts were injected with repair template (Fig. S1). Chimeras were identified and bred to germline confirm transmission. F1 mice from one of the chimeras was bred to C57/BL6j and the line was crossed with new C57/BL6j mice from JAX for three generations before crossing to the other two strains noted in Figure 1G.

### Rats

Timed pregnant Sprague-Dawley rats (E16.5) were purchased from Charles River Laboratories. Upon arrival, animals were housed in a temperature- and humidity-controlled facility with a 12-hour light/dark cycle and ad libitum access to food and water. On postnatal day 0 (P0), litters were collected, and pups were used for culture experiments as described below. All procedures were conducted in accordance with institutional animal care guidelines and approved protocols.

### Primary Cell Culture Protocol for HTS Screening

Forebrains from mice with desired genotype (Fig. 1G) were dissected from post-natal day 0 (PND0) mouse pups to isolate primary cortical neurons in dissection media (culture grade H_2_O (Fisher Scientific: SH3052902), 10% 10x HBSS without Ca^2+^ and Mg^2+^ (Invitrogen: 14185052), 2% HEPES (Invitrogen: 15630080), 1% pyruvate (Invitrogen: 11360070), 1% Glucose solution (Thermo: A2494001), and 0.02% Gentamicin (Invitrogen: 15710064). The cortices were placed in a digestion solution containing dissection media and 20 active units/mL of papain (Worthington: LS003124) for 30 minutes at 37°C. Tissues were washed and triturated in plating medium consisting of Neurobasal (Invitrogen: 21103049) containing 5% heat inactivated FBS, (Invitrogen: 10082139), 2% Glutamax-I, (Invitrogen: 35050061), and 0.02% Gentamicin (Invitrogen: 15710064). Cells were then centrifuged for 5 minutes at 800g and resuspended in plating medium at 300 µL per brain. Cell suspension was then diluted into Feeding medium consisting of Neurobasal-A (Invitrogen: 10888022), 2% Glutamax-I, and 0.02% Gentamicin, 2% B-27 supplement (Invitrogen: 17504044), 10 µM 5-fluoro-2′-deoxyuridine (FUDR) to suppress the proliferation of glia, and 30,000 viral particles/cell of pENN.AAV.hSyn.Cre.WPRE.hGH (AAV9) (Addgene: 105553-AAV9) to induce haploinsufficiency. Using a BioTek EL406 microplate washer dispenser (Agilent Technologies), cells were dispensed into 384-well plates pre-coated with poly-D-lysine (PDL) (Aurora ABE2-01200B-PDL) at 10,000 cells in 80 µL/ well and placed in 37°C incubator. A solution of 1% agarose was placed in the evaporation border wells prior to plating to minimize edge effects. At 7 days in vitro (DIV7), 50% of the conditioned media was replaced with fresh feeding media, and cultures were maintained undisturbed until assayed (usually DIV14).

### HTS-compatible Dual-Luciferase Reporter (DLR) Assay

Neuronal culture plates were assayed (usually at DIV 14) using the Promega Nano-Glo® HiBiT Dual-Luciferase® Reporter Assay System (DLR) (Promega: N1620). Frozen reagents ONE-Glo™ EX Luciferase assay buffer and NanoDLR™ Stop & Glo® Buffer were thawed overnight at 4°C. ONE-Glo™ EX Luciferase Assay Substrate was then resuspended in ONE-Glo™ EX Luciferase assay buffer. All buffers were then equilibrated to room temperature before use. LgBIT protein was diluted 1:100 into ONE-Glo™ EX Luciferase Assay Reagent and NanoDLR™ Stop & Glo® Substrate was diluted 1:100 into NanoDLR™ Stop & Glo® Buffer. 60 µL of culture media from each well of the assay plates was removed using a BioTeK ELx405 (Agilent Technologies) for screen 1 anda BioTek EL406 for screen 2. 10 µL of ONE-Glo™ EX Luciferase Assay Reagent with LgBIT protein was added to each well of the 384-well plate using a BioTeK ELx405 for screen 1 and a Certus Flex liquid dispenser (Trajan Scientific and Medical) for screen 2. Plates were then shaken at 1500 rpm for 10 minutes. The plates were then measured for firefly luciferase (*f*LUC) luminescence using an EnVision plate reader (Perkin Elmer) for screen 1 and a CLARIOstar Plus Microplate Reader (BMG Labtech) for screen 2.10 µL of NanoDLR™ Stop & Glo® Reagent was added to each well of the 384-well plate and shaken at 1500 rpm for 10 minutes. The plates were then measured for NanoBiT (*n*BIT) luminescence on the same reader.

### Compound Administration

#### Primary Screening

Library compounds were administered to neuronal cultured plates on DIV 12 (screen 1) using at 100 nL 384-array pintool (V&P Scientific) or DIV 10 (screen 2) using a 25nL 384-array pin tool (V&P Scientific). The final concentration of the compounds was 3.125 µM or 12.5 µM for screen 1 and 3.125 µM for screen 2 in 80 µL of culture medium. The pin tool was cleaned between each plate by sonicating in water, submerging in DMSO, isopropanol, and methanol, and then dried using house air.

#### Validation pipeline (*Figure 4*)

Neuronal cultures were generated as described above with one modification – feeding occurred every 3-4 days. On feeding days, compounds were readministered using two methods: 50% media exchange with subsequent pinning or by diluting compounds into feeding media and using a Certus Flex dispenser (Trajan Scientific and Medical) to administer media and compound. Pinning compounds requires 100X more compound than feeding, thus the method was chosen based on experimental needs and compound availability.

### Dot Blot Assay

A lysis buffer was prepared containing 2% SDS, 2 mM TCEP, 10% ethylene glycol, 50 mM Sodium Borate, 500 µM 3-(2-Furoyl)quinoline-2-Carboxaldehyde (FQ) (VWR: 102987-910), and 500 µM Mandelonitrile (Sigma:116025) dissolved into water. Using the BioTek EL406, 384-well assay plates containing cortical neurons were washed with 60 µL of PBS three times and then liquid was completely removed using centrifugation. 20 µL of lysis buffer was added to each well using the BioTek EL406 and then shaken at 800 rpm for 10 minutes. The plate was then heated at 75°C using an Envirogenie Incubator (Scientific Industries) for 20 minutes and then cooled at room temperature for 30 minutes. The plate was centrifuged at 3000 g for 3 minutes and shaken for 2 minutes at 400 rpm. Using a pintool array with 384 channels (100nL/channel; V&P Scientific) lysate was pinned on to a 0.2 µm pore size nitrocellulose membrane (Sigma: GE10600004), dried for 1 minute, and placed into a container with 1X TBS-T. The membrane was then blocked with 1% BSA-TBS-T for 1 hour. Antibodies (*see below)* were then diluted into 1% BSA-TBS-T at appropriate concentration and incubated overnight at 4°C on a platform rocker. The membranes were washed 3X with TBS-T for 10 minutes and then incubated with HRP-conjugated secondary antibodies *(see below)*. The membranes were washed 3X with TBS-T and once with TBS. Membranes were then imaged for total protein using BioRad ChemiDoc imaging system using the Stain Free Blot setting. 15 mL of SuperSignal West Pico Plus Chemiluminescent substrate (Thermo:34580) was added to the membrane and incubated for 2 hours on a platform rocker, followed by addition of 2 mL SuperSignal West Femto Maximum Sensitivity substrate (Thermo: 34096), and incubated for 5 minutes, and imaged for chemiluminescence.

### SDS-PAGE and Immunoblotting

Primary neuronal cultures were prepared as described above (validation protocol) from *Syngap1* heterozygous cRescue mice and plated within PDL coated 24-well plates at 250,000 cells / well. FUDR was added at DIV 3 instead of at plating. Cultures were maintained by 50% media exchanges every 3-4 days. On DIV 14, plates were washed with PBS twice and proteins were extracted by sonication in a buffer consisting of 2% SDS, 50 mM Sodium Borate, 1X Halt Protease and Phosphatase inhibitors. Sample protein concentrations were measured using Pierce BCA Protein Assay Kit (Thermo: 23225) and adjusted to normalize protein content. 10µg of protein per sample was loaded and separated by SDS-PAGE on 10% Criterion TGX Stain-Free gels (BioRad: 5678035) and then transferred to low fluorescence PVDF membranes (45 µm pore size) (Amersham: GE10600004) with the Trans-Blot Turbo System (BioRad). Membranes were imaged for total protein using BioRad ChemiDoc imaging system, blocked with 1% BSA-TBS-T for 1 hour, and then probed with primary antibodies at 4°C overnight. Membranes were washed 3X with TBS-T, incubated with secondary antibodies, washed, and imaged for chemiluminescence. Direct *n*BIT luciferase signal was also measured using the Nano-Glo® HiBiT Blotting System (Promega: N2410). Following chemiluminescence detection, 30% H_2_O_2_ was added to the membrane to quench the chemiluminescence and washed 3X with TBS-T. LgBiT protein was added in Nano-Glo Blotting buffer, incubated overnight at 4°C, and Nano-Glo® Luciferase assay substrate was diluted 500-fold into the Nano-Glo blotting buffer. The membrane was incubated for 5 minutes and then imaged for chemiluminescence.

#### Primary Antibodies

Pan-SynGAP antibody (1:1000) – Cell Signaling #5539

SynGAP-α2 antibody (1:1000) - Cell Signaling #56927

SynGAP-α1 antibody (50 ng/mL) – Cell Signaling Test sample (VSP-137655); Rabbit mAb #34124 SynGAP-β antibody (50 ng/mL) - Cell Signaling Test sample (VSP-143511); Rabbit mAb #28580 ANTI-FLAG M2 antibody (1:1000) – Sigma #F1804

#### Secondary Antibodies

Anti-mouse IgG HRP Conjugate (1:2500) – Promega W402B Anti-rabbit IgG HRP Conjugate (1:2500) – Promega W401B

### Cell-Free *n*BIT (Counter-screen) Assay

A counter screening assay was developed to determine to what extent small molecules directly regulated *n*BIT activity. Starting with the Nano-Glo® HiBIT Extracellular Detection System (Promega: N2420), a HiBIT control protein (Promega: N3010) was diluted to 100 pM in buffer containing 0.1% BSA-PBS and 10 µL was dispensed into black opaque 384-well assay plates (Greiner: 781900). Compounds were pinned into the plate and shaken at 1500 rpm for 1 minute. 10 µL of Nano-Glo® HiBIT Extracellular buffer with 1:50 Nano-Glo® HiBIT Extracellular substrate and 1:100 LgBIT protein was dispensed into the plate, shaken for 1 minute, centrifuged at 100 g for 1 minute, and incubated for 10 minutes. The plate was then measured for luminescence using the CLARIOstar Plus plate reader.

### Screening Libraries

Both custom designed and pre-selected screening libraries were obtained from Enamine (Ukraine) and ChemBridge (San Diego). Libraries were delivered in 384-well plates and each well contained ∼30uL of compound (10mM). Compounds were absent in columns 1, 2, 23, 24 to accommodate controls. Assay plates used in the HTS-style screen were pinned directly from the library compound plates. Controls were pinned separately.

### Electrophysiology and mEPSC Analysis

Primary neuronal cultures were prepared as described above from *WT* and constitutive heterozygous *Syngap1* KO mice and plated onto PDL coated glass coverslips (Neuvitro: GG-12-pdl) in 24-well plates at 250,000 cells / well. FUDR was added at DIV 3 instead of at plating. Cultures were treated with vehicle (DMSO) or SR-1815 (1.5 µM) on DIV 0, 3, 7, and 10 using a 50% media exchange. Whole-cell patch clamp recordings were conducted from forebrain cultures between DIV 13-15. Putative excitatory neurons were visually identified using infrared DIC optics. Recordings were made using boroscilicate glass pipettes (3-6 MΩ; 0.86 mm inner diameter; 1.5 mm outer diameter; Harvard Apparatus) made using a P-97 pipette puller (Sutter Instruments). All signals were amplified using Multiclamp 700B amplifier (Molecular Devices), filtered at 2.4 KHz, digitized at 10 KHz and stored on a personal computer for off-line analysis. Analogue to digital conversion was performed using a Digidata 1400A system (Molecular Devices). Data acquisition and analysis was performed using pClamp 10.2/11.2 software (Molecular devices), along with Minianalysis software (Synaptosoft) for semi-automated mEPSC event detection. To isolate AMPA-mediated mEPSCs, cultures were continuously perfused with artificial cerebrospinal fluid (aCSF), composed of (in mM): 150 NaCl, 3.1 KCl, 2 CaCl_2_, 1 MgCl_2_, 10

HEPES, 10 Glucose, 0.05 D-2-amino-5-phosphonovalerate, 0.001 tetrodotoxin and 0.1 picrotoxin. Osmolarity was adjusted to 305-310 mOsm and pH adjusted to 7.3-7.4. The internal solution consisted of (in mM): 135 Cs-methanesulfonate, 10 CsCl, 10 HEPES, 5 EGTA 2 MgCl_2_, 4 Mg-ATP, and 0.1 Na-GTP. The internal solution was adjusted to pH 7.3 and to 290-295 mOsm. Following the establishment of whole-cell configuration, passive membrane properties were monitored throughout the experiment. Cells with access resistance > 30 MΩ or unstable (>20 % change) were discarded from analysis. A minimum of 100 and a maximum of 200 events were collected from each neuron, while 2-4 neurons were patched per culture. The amplitude threshold for event detection was set to √RMS x3 (typically ∼4 pA). Data acquisition and analysis were performed by the same experimenter blinded to both treatment and genotype. Data from each group was averaged and statistical significance was determined using 2-way ANOVA with Fisher’s LSD posthoc test. Data are expressed as mean ± SEM.

### Calcium Imaging and Analysis

#### Imaging

Primary neuronal cultures were prepared as described above and plated into PDL coated 384-well plates at 15,000 cells / well. FUDR was added at DIV 3 instead of at plating. Cultures were maintained by 50% media exchanges and treated with vehicle (DMSO) or SR-1815 on DIV 0, 3, 7, and 10. On DIV 3 cultures were transduced with AAV9 vectors expressing pGP-AAV-syn-FLEX-jGCaMP8f-WPPRE (AAV9) (Addgene: 162379-AAV9) (MOI=300,000 vp/cell) and pENN.AAV.hSyn.Cre.WPRE.hGH (AAV9) (Addgene: 105553-AAV9) (MOI=10,000 vp/cell). On DIV 14, imaging was performed on an InCell Analyzer 6000. Image series were acquired through a 20x Objective (Nikon 20X/0.45, Plan Fluor, ELWD, Corr Collar 0-2.0, CFI/60) at a rate of ∼10Hz using a 478nM laser for sensor excitation. Neuronal activity was analyzed from time-lapse fluorescence microscopy images. Images were first segmented using the Cellpose deep learning model to identify individual neurons. The average fluorescence intensity within each segmented region was extracted over time. These signals were then detrended and normalized to obtain ΔF/F values. Spikes were detected in the ΔF/F traces by setting a threshold based on the baseline noise level. Spike frequency was calculated for each neuron. The results were aggregated across all analyzed images, combining spike metrics with segmentation properties and experimental metadata (e.g., drug treatment, GCaMP type).

### Hit Identification for Screen 1

Each compound plate was tested in duplicate on two plates containing cultured neurons at DIV 12. At DIV 14, the DLR assay was performed and yielded two *f*LUC and two *n*BIT reads per compound plate tested. To identify hits from each compound plate, the median *f*LUC and *n*BIT values of the negative controls (DMSO only) were calculated for each plate. This value was used to normalize each plate and then each duplicate read was averaged for both *f*LUC and *n*BIT. Using the averaged normalized data, the standard deviation and mean of the negative controls was used to calculate a z-score for each data point. The data was plotted on a 2D scatter plot with the z-score of the normalized data for *f*LUC on the x-axis and the z-score of the normalized data for *n*BIT on the y-axis. To identify hit compounds, vertical threshold lines were drawn at 3 X standard deviation (SD) left and right for *f*LUC and horizontal lines above and below for *n*BIT (z-score = ± 3). Compounds with *f*LUC values within 3X SD and *n*BIT values above 3X SD were identified as potential upregulators, whereas those with nBIT values below 3X SD were considered down-regulators. To ensure reproducibility, variability filtering was applied. The coefficient of variation (CV) was calculated for each compound’s *f*LUC and *n*BIT replicate reads. Only compounds with <10% variability for both *f*LUC and *n*BIT were retained as final hits. 2D scatter plots were color coded for each category: negative control (green), compound field (gray), and hits (blue).

### Development of Hit Detection Algorithm for Screen 2

All experiments are performed in duplicate. This yields two *f*LUC (*f*_1_, *f*_2_) and two *n*BIT (*b*_1_, *b*_2_) reads per experiment. The 1^st^, 2^nd^, 23^th^, and 24^th^ datapoints from rows “A”, “B”, “O”, and “P” are excluded from the analysis as these are often affected by edge effects. First, a logarithmic transformation is applied to all raw luminescence intensities (log_2_*f*_1_, log_2_*f*_2_, log_2_*b*_1_, log_2_*b*_2_).

Second, a Mahalanobis-distance^50^ based filter is applied to the negative control data, allowing us to correctly estimate the distance of each negative control datapoint from the center of mass of the distribution in a direction dependent manner, to remove obvious outliers.

The Mahalanobis-distance *DM_i_* is calculated as follows for all the 𝑛 observations in the negative control:

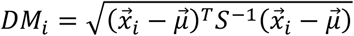

where 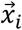 is a column vector representing the *i* ^th^ observation:

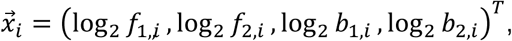

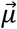 is a column vector representing the mean of the negative control:

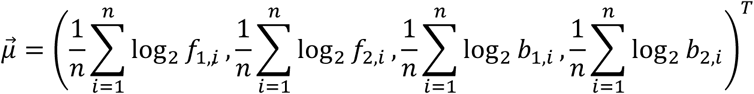

𝑆^−1^ is the inverse of the covariance matrix, and ^𝑇^denotes transposition. Any datapoint with a Mahalanobis distance larger than 3 is considered an outlier.

Next, data reduction is done by calculating the mean of the two reads for all datapoints:

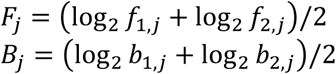

Where 𝐹_𝑗_ and 𝐵_𝑗_ are the reduced *f*LUC and *n*BIT signals of the *j* ^th^ observation, respectively.

The reduced data is then normalized by subtracting the mean of the outlier-filtered negative control group from each datapoint:

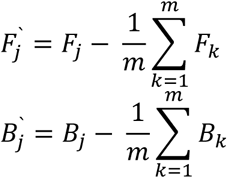

Where 𝐹_𝑗_ and 𝐵_𝑗_ are the normalized *f*LUC and *n*BIT signals of the *j* ^th^ observation, 𝐹_𝑘_ and 𝐵_𝑘_ are the reduced firefly and nanoBIT signals of the *k*^th^ observation among *m* total observations in the outlier filtered negative control group, respectively.

This transformation shifts the coordinates of the center of the negative control ellipsoid to (0,0) in a two-dimensional double-logarithmic scatter plot (Fig. S3A).

The effect size for each data point is calculated as the difference of the observed and expected *n*BIT signals (Fig. S3B). The observed signal is simply the normalized *n*BIT signal (as defined above), while the expected *n*BIT signal equals the normalized firefly signal:

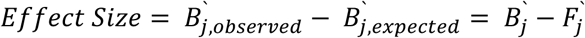

This relationship is the consequence of the linear correlation between cell density and protein levels in a sample. Any relatively small change in the cell density that does not result in a change of the “fundamental” biological processes (e.g. protein expression levels) in the sample must be followed by a similar change in both the *f*LUC and *n*BIT signals. Interestingly, one can observe this correlation in positive and negative control data (Fig. S3C), even if these controls are not designed to detect this phenomenon. In the controls from screening, only the random experimental variability (e.g. dispensing of a cell suspension) results in different cell densities.

Due to random experimental variability and the above-described relationship, compounds with the same effect size are expected to be scattered along lines with a slope = 1 in double logarithmic scatter plots (Fig. S3A-B). To identify hits in a screening experiment, threshold lines with a slope of 1 are required (Fig. S3A-B). Compounds showing significant firefly effects relative to the negative control (e.g. due to toxicity, induction of non-specific protein expression changes, inhibition, activation or stabilization of firefly luciferase^51^) should not be considered as hits. To identify such compounds, two vertical threshold lines are used at 0 ± 3 x standard deviation of the negative control (Fig. S3A-B*, vertical orange lines*).

Last, the “non-repeatedness” defined as (log_2_*f*_2_ - log_2_*f*_1_) - (log_2_*b*_2_ - log_2_*b*_1_) is calculated for all datapoints and reflects how much the two parallel experiments do not replicate each other. If the non-repeatedness value is above X then the compound is not selected as a hit. (Fig. S3F).

### Automated Dot Blot Analysis

An Image J macro was developed to automate the dot blot image analysis process. The analysis involves fitting a grid to the image representing the microplate from which the samples were transferred to the membrane (Fig. S5A-D). The distance between the center of any two immediate neighbors in this array (in pixel units) is theoretically constant and represents the distance of neighboring wells in the plate. In order to find dots representing the real samples, shape-based sample identification is performed by running a sequence of operations. First, edge detection is performed and the resulting image is transformed into a binary (black and white) image (Fig. S5B). This is followed by local maxima identification, which gives a set of experimental coordinates representing the center of samples. To remove high intensity artifacts, the “flood fill tool” is used followed by selecting the whole object (region of interest), which includes the area surrounded by the circular edge and the edge itself. Real dots on the membrane give selections with very similar area and almost perfectly circular shape. Artifacts, however, typically give selections with much smaller area and irregular shapes (Fig. S5B). Therefore, the identified objects are filtered by size and median width, median height and median circumference are calculated. These statistics are used as shape-based filters to remove any remaining artifacts from the list.

Next, with a set of coordinates representing real samples only, the optimal distance between the closest neighbors is determined (in pixel units) by generating a histogram from the calculated Euclidian distance between the center of each spot and the center of all other identified spots in the image using the following formula:

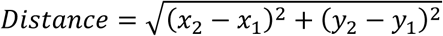

Where (*x_1_*, *y_1_*) and (*x_2_*, *y_2_*) are the coordinates of the 2 spots in pixel units (Fig. S5C). A theoretical distance distribution is also calculated (Fig. S5C) by assuming unit distance (= 1) between the closest neighbors in an idealized grid representing the plate. The optimal (experimental) distance “*d*” is determined by fitting the theoretical histogram to the experimental histogram. This is done by multiplying the peak positions in the theoretical histogram by a variable “*d”* and optimizing the value of “*d*” such that the peak positions in the theoretical histogram overlap with the peak positions in the experimental histogram. The position of the peaks is compared by defining a zone for each peak in the theoretical distance histogram such that the boundaries of the zone fall half-way in between the neighboring peaks. The sum of the distances of each theoretical peak and the experimental peaks within its zone are calculated. The global minimum of this penalty function gives the optimal distance (*d*) in pixel units.

Next, the orientation of the membrane within the image needs to be determined. Each sample has a maximum of four closest neighbors allowing a displacement vector to be defined pointing to each of these neighbors with coordinates (*x_2_*-*x_1_*, *y_2_*-*y_1_*), where *x_2_*-*x_1_* and *y_2_*-*y_1_*are the displacement components along the “x” and “y” axes, respectively, (*x_1_*, *y_1_*) are the coordinates of the sample, and (*x_2_*, *y_2_*) are the coordinates of the neighbor in pixel units (Fig. S5D). Plotting these vectors for all samples yields four distinct populations around the origin. The distance between the center of each of these populations and the origin is expected to be equal to the optimal distance between the closest neighbors (see previous step). Relative to any arbitrarily chosen one of the four vectors pointing to the center of these populations, the other three are rotated by 90°, 180° and 270°. Therefore, for each datapoint, we calculate all possible rotations (0°, 90°, 180° and 270°) and chose the one with the angle closest to zero in all cases. The mean of all the angles represented by the datapoints yields the optimized orientation (*α*).

Next, a relationship between the coordinates of samples (in pixel units) and coordinates in an idealized plate (row, column) needs to be determined (indexing). Image coordinates are transformed by a combination of a translation, a rotation by *α*, and a division with the optimal distance (*d*) determined above:

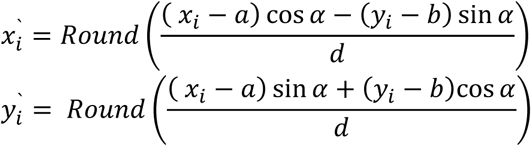

Where *x_i_’* and *y_i_’* are the transformed coordinates, *x_i_* and *y_i_* are the image coordinates of the i^th^ sample, *a* and *b* are the image coordinates of one randomly selected sample, and “Round()” denotes rounding to the nearest integer. The transformed coordinates are then translated to set the smallest *x_i_’* and *y_i_’*values to (0,0).

The indices *x_i_’* and *y_i_’* are used to generate a set of idealized coordinates by using the optimal distance and reversing the transformations. The sum of Euclidian distances between each sample and the corresponding location in the idealized grid is calculated and used as a penalty function in coordinate refinement. A series of translations is performed along the x and y axes until the global minima of the penalty function (a 2D surface) is found (Fig. S6A). At this point, the image coordinates of sample “A01” (column = 1, row =1) define the optimal location of the grid in the original image (x_A01_, y_A01_; see Fig. S5A).

Next, missing samples are identified in the list of indices, and the above defined transformations are performed in reverse order to obtain the corresponding coordinates of the grid.

In the last step, sample integration is done with local background correction (Fig. S6B). Pixels around each grid point in the unmodified image are selected within a circle covering the entire spot to calculate a “raw” integral. This raw integral needs to be corrected for background, which can be estimated from the integral of the immediate surroundings of the circle. We selected pixels in a larger circle such that the center of both circles is located at the coordinates defined by the grid point, and the area of the large circle is exactly 2 times the area. This gives a radius of *r_L_* = *r*\*√2. The integral within the small circle is a sum of the signal (*S*) and the background (*B*): *I_1_* = *S* + *B*. The integral within the large circle contains the signal (*S*) and twice the background (*B*): *I_2_* = *S* + 2*B*. Thus, the background corrected signal can be calculated as *S* = 2*I_1_* – *I_2_*. Although the radius can be easily optimized by testing all possible *r* values between 1 pixel and *d*/(2*√2) pixels we typically chose *r* = *d*/(2*√2) to avoid high variability due to partially lost signal in areas of local membrane distortions and to minimize the effect of overestimated background due to high intensity local artifacts.

## Acknowledgements

We are grateful to the entities that provided sponsored research funding that supported the development of this platform (National Institute for Mental Health - U01MH136567 and R01MH113648; Praxis Precision Medicines - UFAS-A00002). A custom chemical library was curated by Andy Jennings and shared with the Rumbaugh Laboratory by Praxis Precision Medicines. Jordan Hirschfeld of Cell Signaling Technology (CST) developed and shared two SynGAP isoform-specific antibodies *(see methods)*. Lina Deluca handled all compound management activities.

## Supplemental Figures

**Supplementary Figure 1.**
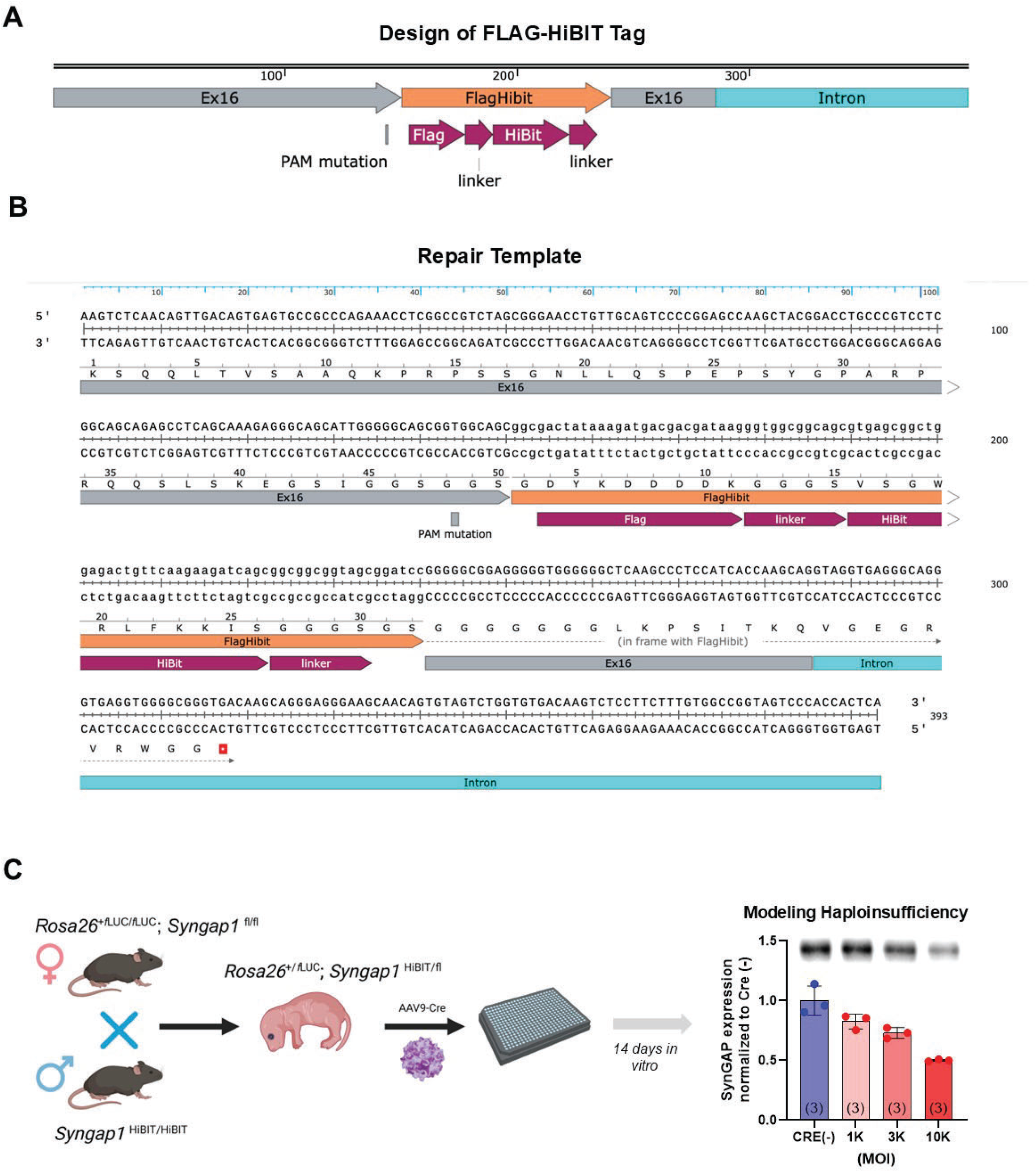
Design of Syngap1-HiBIT knock-in mouse line. **A)** Design of the HiBIT tag to be inserted into the murine *Syngap1* locus using CRISPR. **B)** Sequence of the repair template and location of the PAM site. **C)** Demonstration that application of a viral vector expressing Cre recombinase during culture plating induces *Syngap1* haploinsufficiency in primary mouse neurons destined for high-throughput-style screening. Error bars represent SD.

**Supplementary Figure 2.**
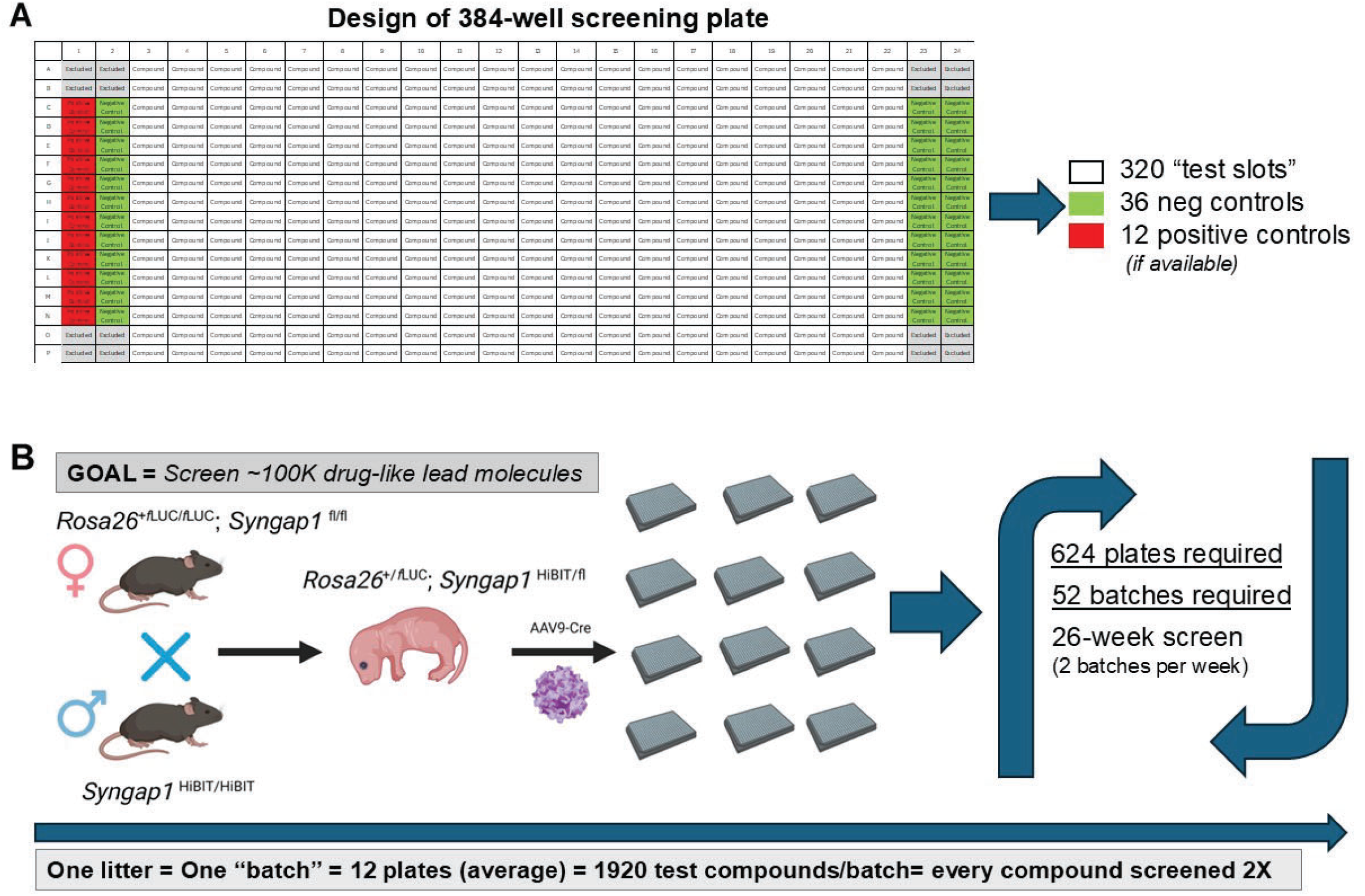
Layout of screening plate and design of 100K compound screen based on this plate design. **A)** Location of negative control (green), positive control (red), and test agent positions (320 compound slots) in each screening plate for Screen 2. Screen 1 negative controls were used in column 1. **B)** One litter of mice can regularly produce up to 16 assay plates (depending on the desired neuronal plating density). To screen 100K compounds in duplicate, 624 plates would be required across ∼52 individual 12-plate batches of primary cortical neurons, which would take ∼6 months to complete when culturing two batches of neurons per week.

**Supplemental Figure 3.**
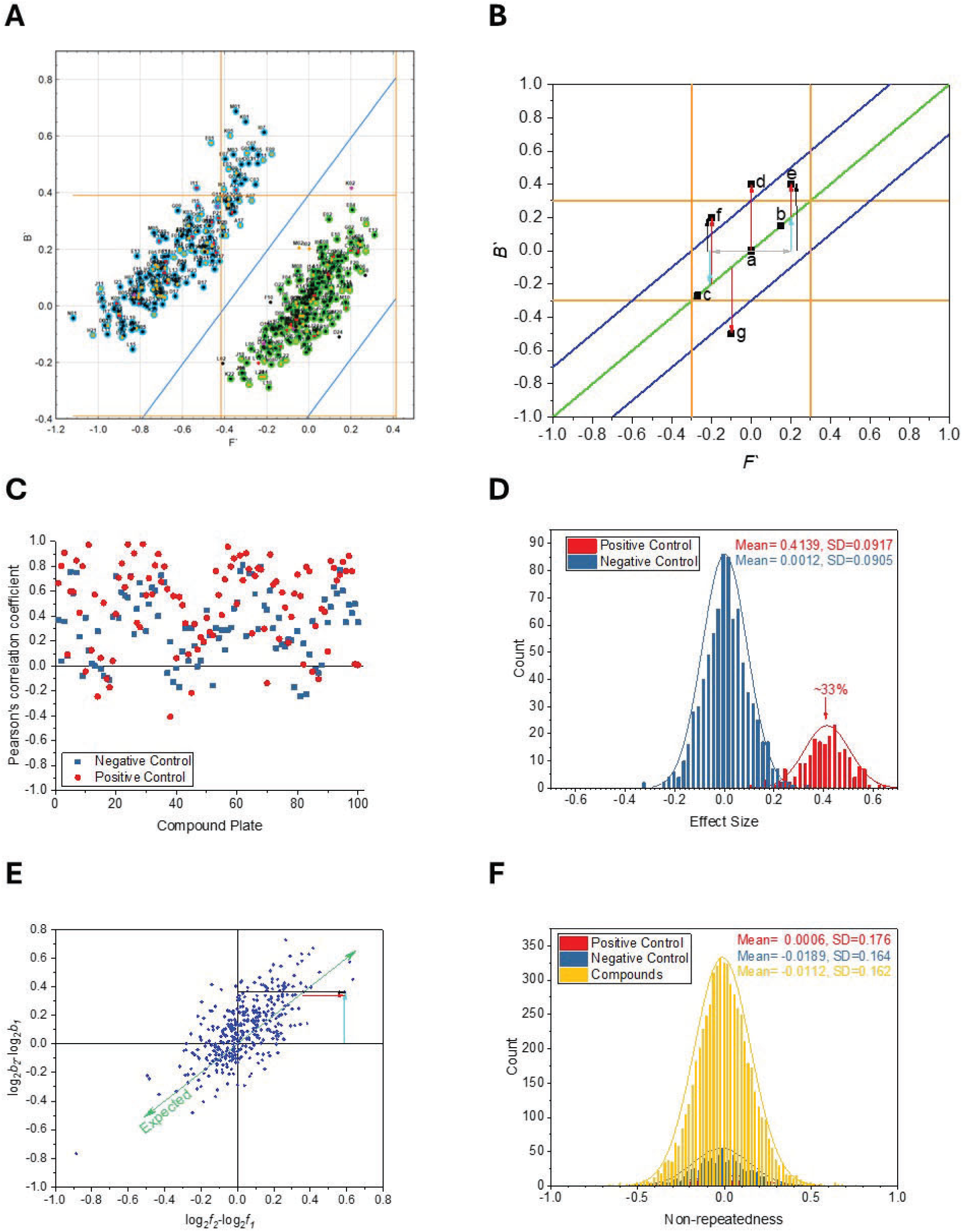
Screening data exploration, analysis, and subsequent development of hit-detection algorithms. **A)** Double logarithmic 2D-Scatter plot showing half-plate negative and half-plate positive control normalized data (highlighted by green and blue ovals, respectively). Orange lines represent 0±3 x standard deviation of the negative control. The elongated shape of the ellipsoids indicates that the two signals (*F’*, *B’*) are correlated in both controls (see also panel C). Therefore, horizontal threshold lines must be “tilted” (blue, slope =1) to separate the controls correctly. Inner circles are color-coded to represent the “non-repeatedness” (black ≤ 1xStDev, 1xStDev < orange ≤ 2xStDev, 2xStDev < red ≤ 3xStDev, 3xStDev < violet). **B)** Samples with no effect in double logarithmic plots (“a”, “b”, and “c”) are expected to show *B’* = *F’*. The green line with a slope = 1 and intercept = 0 symbolizes this relationship. The observed signal *B’_j_* for the j^th^ sample (black arrows, only two examples shown for clarity) needs to be corrected with the expected signal (cyan arrows), which equals the observed *F’_j_* (gray arrows) to estimate the effect size (red arrows). Note that for the upregulators “d” and “e”, the effect size is different, while for “d” and “f” are equal. Also, sample “g” is a downregulator showing an effect size with the same magnitude but an opposite sign. Samples with the same absolute effect size (“d”, “f”, and “g”) are correctly identified as hits by the blue threshold lines. **C)** Pearson’s correlation coefficients for positive and negative controls (100 plates) are typically greater than zero, thus indicating positive linear correlation. **D)** Effect sizes for both positive and negative controls show a symmetrical distribution due to the logarithmic transformation applied to the data. Note, the mean of the negative control is zero, while the positive control is shifted to the left, representing a mean effect size (upregulation) of ∼0.41 (equivalent to ∼33%). Data was collected in a pilot optimization study of 29 plates (negative control: 42 wells/plate, positive control: 14 wells/plate). **E)** The “Non-repeatedness” (red arrow) is a measure of the unexpected component of the difference between log_2_*f*_2_ -log_2_*f*_1_ (black arrow) and log_2_*b*_2_ - log_2_*b*_1_ (cyan arrow), which are the differences of the two log_2_-transformed firefly (*f*_1_, *f*_2_) and nanoBIT reads (*b*_1_, *b*_2_), respectively. **F)** Non-repeatedness data of the control groups and compound fields from 29 plates show symmetric distributions centered at ∼0 and very similar standard deviations. Thus, the standard deviation of the negative control is used to estimate data quality (see the color coding in panel A).

**Supplementary Figure 4.**
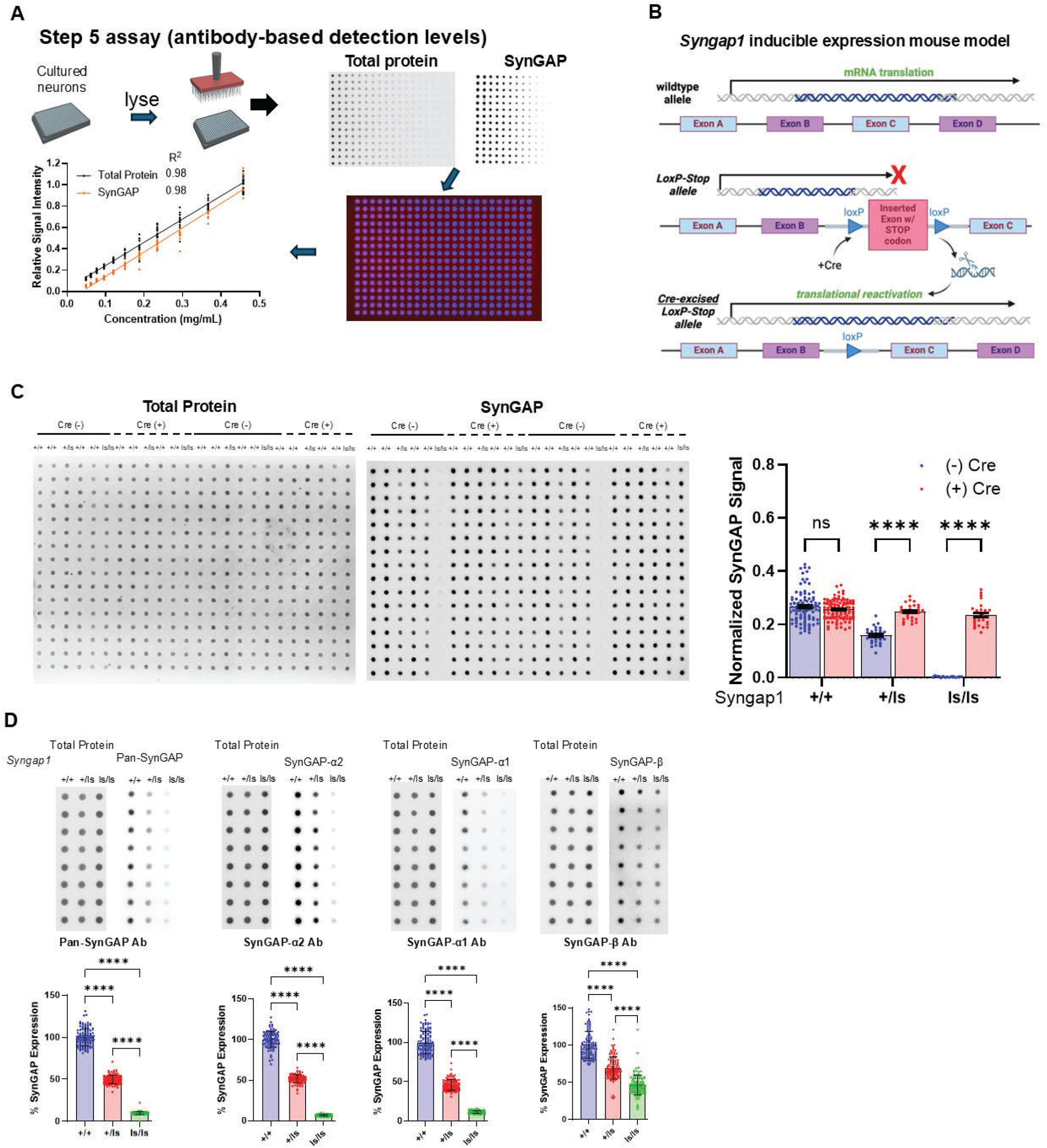
Development and validation of a scalable protein Dot Blot assay to measure endogenous SynGAP protein. **A)** Workflow for Dot Blot assay. Cultured neurons are lysed and pinned onto membrane. Linearity of Dot Blot signals for total and SynGAP protein levels in neurons, simple linear regression for both total protein and SynGAP, p=<1.0*^e^*15, n=14 per concentration. **B)** Cartoon demonstrating features of the Cre-conditional *Syngap1* cRescue mouse line**. C)** Biological validation of Dot Blot assay using neurons cultured from conditional *Syngap1*-rescue mouse line. *“ls”* is the targeted conditional *Syngap1* “rescue” allele. Images of Dot blot assay showing total protein (*left*) and SynGAP (*middle*), with quantification of Dot blot (*right*; 2-way ANOVA with Tukey’s post hoc; Genotype: F (2,316) = 268.2, p=<1.0*^e^*15, Cre: F (1, 316) = 343.6, p=<1.0*^e^*15, Interaction: F (2,316) = 194.1, p=<1.0*^e^*15). For (-) Cre *Syngap1^+/+^* n=98, *Syngap1^+/ls^* n=28, *Syngap1^ls/ls^* n=98; for (+) Cre *Syngap1^+/+^* n=112, *Syngap1^+/ls^* n=28, *Syngap1^s/ls^* n=28. Error bars represent SEM. Background subtraction was performed by subtracting average *Syngap1^ls/ls^* for each data point. **D)** Dot blot assay using total SynGAP and SynGAP isoform specific antibodies from *Syngap1^+/+^, Syngap1^+/ls^*, and *Syngap1^ls/ls^* neurons. % SynGAP expression was normalized to total protein and normalized to *Syngap1^+/+^*.One-way ANOVA with Tukey’s post hoc tests were used for all analyses. Pan-SynGAP: F (2, 333) = 5020, p=<1.0*^e^*15. SynGAP-α2: F (2, 290) = 5096, p=<1.0*^e^*15. SynGAP-α1: F (2, 333) = 2682, p=<1.0*^e^*15. SynGAP-β: F (2, 333) = 328.6, p=<1.0*^e^*15. n=112 for each genotype and antibody tested. Error bars represent SD.

**Supplementary Figure 5.**
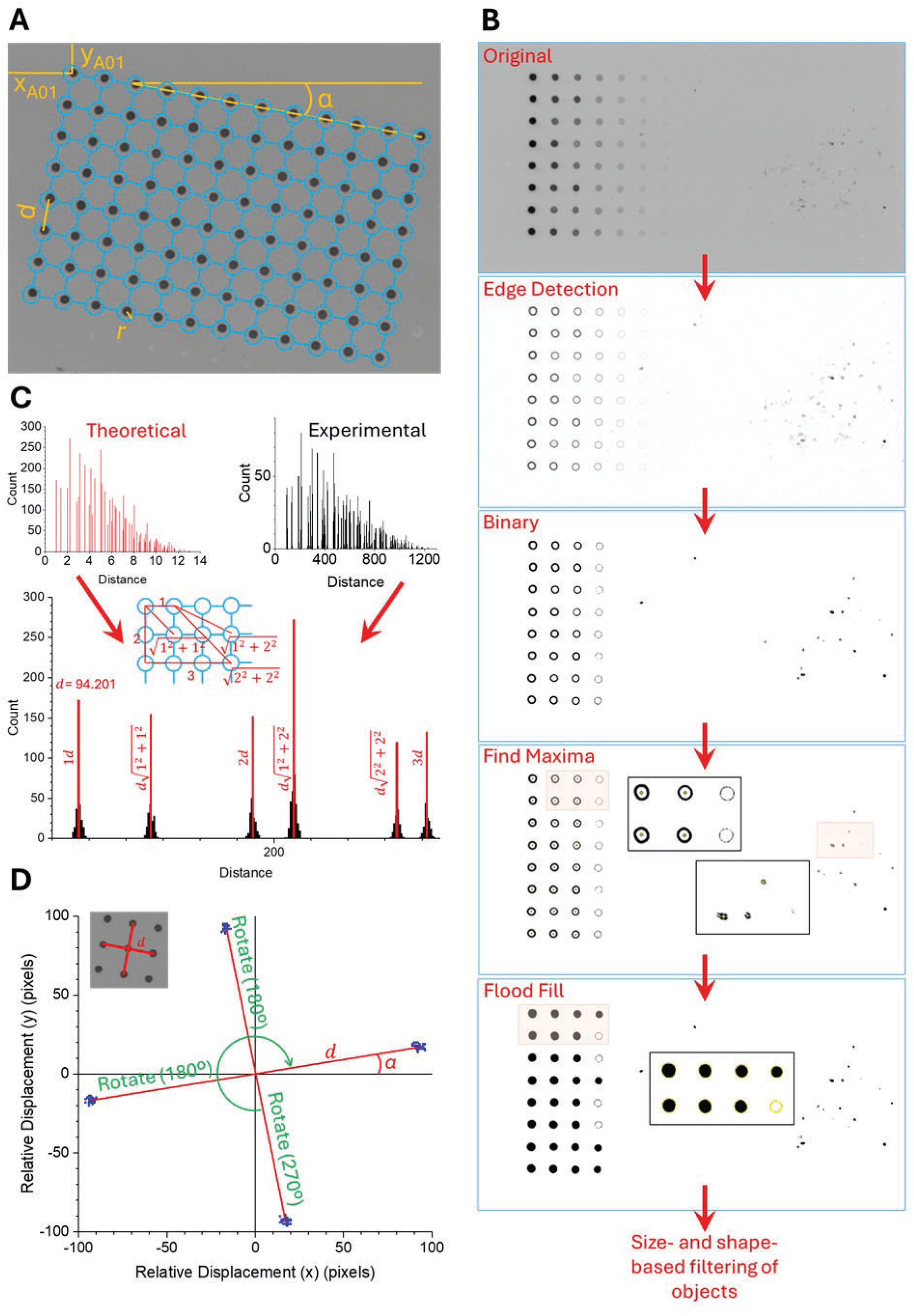
Automated analysis of Dot Blots. **A)** Fitting an ideal grid representing a microplate to a Dot Blot involves the optimization of 5 parameters: “x” and “y” coordinates of the “northwest” corner of the grid (*x_A01_*, *y_A01_*), distance between the closest neighbors (*d*), orientation of the membrane relative to the x axis of the image (*α*), and the radius of the circular selection of pixels where the integral is calculated (*r*). Only 96 wells of the 384-well plate were used here for simplicity. **B)** An image showing a Dot Blot calibration experiment with “invisible” (very low intensity) samples and many artifacts is intentionally shown for demonstration purposes. Identification of real samples starts with edge detection, followed by conversion into a binary (black and white) image, maxima identification, flood fill, selection of objects, and finally, size and shape-based filtering. Maxima are shown as yellow dots and used in subsequent steps (after removing artifacts) as experimental coordinates to fit the grid. Highlighted areas are zoomed in and presented as insets. **C)** A theoretical distance histogram (red) is a collection of all possible Euclidean distances measured from every possible location to every other location in an ideal grid with a unit (=1) distance in between the closest neighbors. The experimental distance histogram (black) is a collection of all possible Euclidean distances measured from every experimentally identified sample (coordinates of the center) to every other sample in an image. The inset gives one example for the calculation of each of the first 6 distances (peak positions) in the theoretical histogram. These peak positions are multiplied by the variable “*d*” to “stretch” the histogram until the theoretical and experimental peaks are closest to each other. At this point, *d* equals the optimal distance (94.201 pixels in this example). **D)** The orientation of the membrane (*α*) is determined by calculating all possible displacement vectors (red) pointing from the center of every sample to the center of its closest neighbors. Note that the length of these vectors is ideally the optimal distance “*d*”. The displacement components along the “x” and “y” axes of these vectors give 4 distinct populations (blue). Each datapoint represents an angle. The rotations shown are performed to align these populations such that this angle is closest to zero. The mean of all angles (*α*) is the optimal orientation of the membrane relative to the edge of the image.

**Supplementary Figure 6.**
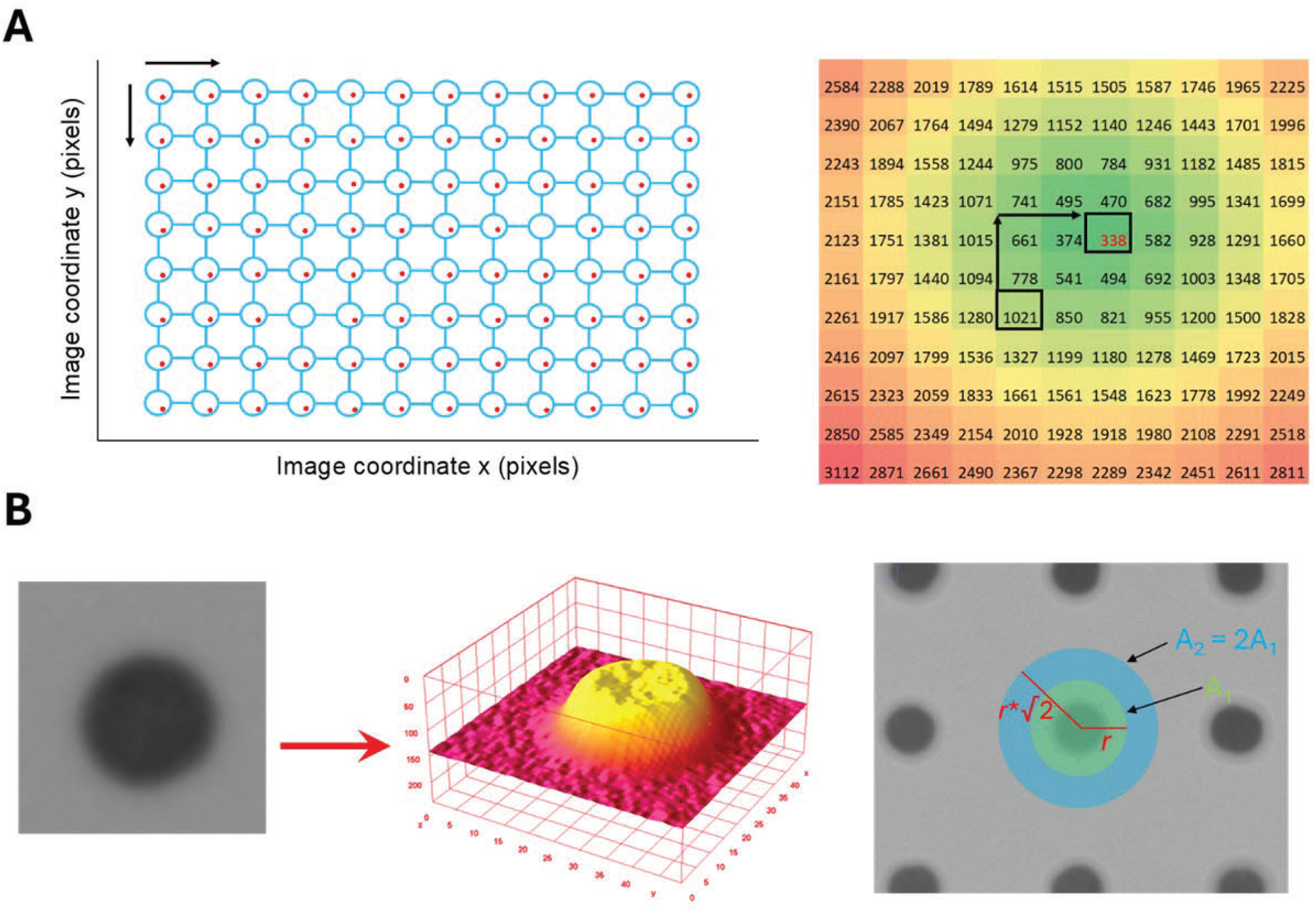
Automated analysis of dot blots (cont’d). **A)** The coordinates of the grid representing an ideal plate (center of blue circles) is aligned to the experimentally determined coordinates (red dots) by minimizing the sum of Euclidian distances between the experimental and the corresponding grid coordinates (penalty function). Black arrows show the direction of the translations performed during the alignment until the global minima of the penalty function (shown as a heatmap) is found. (The algorithm does not calculate the whole surface. It rather “walks” along the surface towards lower values by comparing values of the closest neighborhood only.) At the global minimum of the penalty function, coordinates of grid point “A01” define the “northwest” corner of the grid (*x_A01_*, *y_A01_*). **B)** A sample on the membrane can be represented as a surface. Integration over a circular selection with a radius “*r*” and area “*A_1_*” (green circle) gives the volume under this surface (*I_1_*). Note that the background intensity may contribute significantly to the volume. Therefore, local background correction needs to be performed by calculating a second integral (*I_2_*) over a larger circular selection (blue circle) such that the radius equals *r*\*√2 and thus the area *A_2_*= 2*A_1_*. The background corrected signal is given by 2*I_1_*- *I_2_*.

**Supplementary Figure 7.**
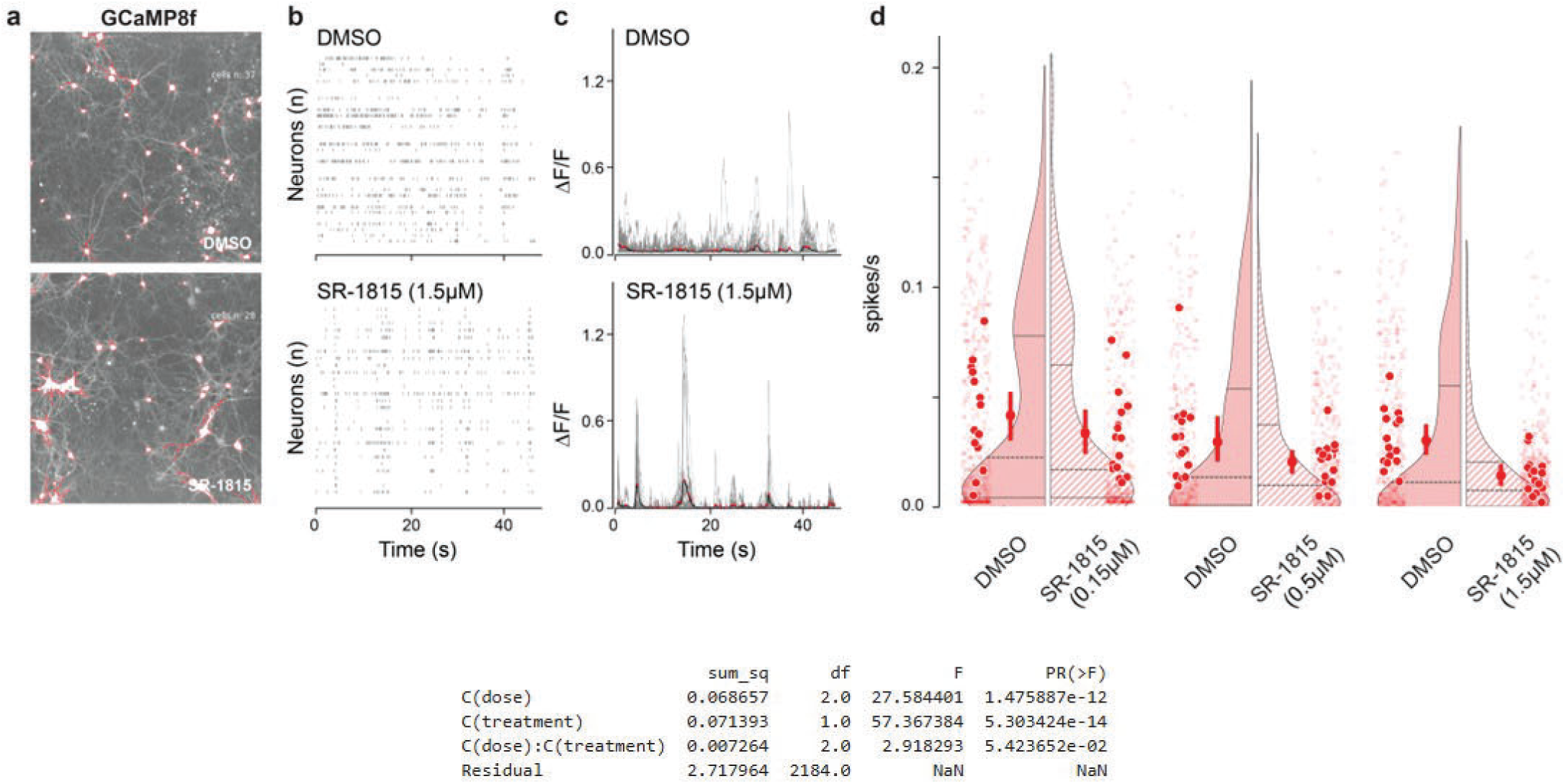
SR-1815 drives a dose-dependent reduction in neuronal activity within *Syngap1* haploinsufficient neurons. **A)** Maximum projection of 2-minute movie of GCaMP dynamics. The red outlines reflect automated cellular segmentation based on maximum intensity projection for vehicle and SR-1815 in culture transduced with GCaMP8 and treated with SR-1815 (1.5 μM). **B)** Spike raster of calcium transients. **C)** ΔF/F of calcium signals from neurons segmented in one field of view. **D)** Spike rate for DMSO and SR-1815 at 3 concentrations (low=1.5 μM, mid=0.5 μM, and high=1.5 μM. Large dots plotted over the smallest dots represent sample average for the plotted measure (n=16 per condition) and average of these averages correspond to the dot with the bar representing the sem. Plot showing the spiking frequency (spikes per seconds) for the different conditions (OLS regression with Tukey HSD post hoc test, Dose: F(2,2184)=27.6, p=1.5e-12, Treatment: F(1,2184)=57.4, p=5.3e-14, Interaction: F(2,2184)=2.91, p=5.4e-2). For DMSO and SR-1815 16 fields were imaged in total from 2 wells for the three different doses for *Syngap1^+/-^*. The total number of segmented neurons were DMSO low, mid, high: 360,310,376 and SR-1815 low, mid, high: 278,413, 453.

